# Fungal zygote survival and ploidy maintenance rely on cell cycle-dependent and independent mating blocks

**DOI:** 10.1101/2020.07.26.222000

**Authors:** Aleksandar Vještica, Melvin Bérard, Gaowen Liu, Laura Merlini, Pedro Junior Nkosi, Sophie G Martin

## Abstract

To ensure genome stability, sexually reproducing organisms require that mating brings together exactly two haploid gametes and that meiosis occurs only in diploid zygotes. In the fission yeast *Schizosaccharomyces pombe*, fertilization triggers the Mei3-Pat1-Mei2 signaling cascade, which represses subsequent mating and initiates meiosis. Here, we establish a degron system to specifically degrade proteins post-fusion and demonstrate that mating blocks not only safeguard zygote ploidy but also prevent lysis caused by aberrant fusion attempts. Using long-term imaging and flow-cytometry approaches, we identify previously unrecognized and independent roles for Mei3 and Mei2 in zygotes. We show that Mei3 promotes premeiotic S-phase independently of Mei2 and that cell cycle progression is both necessary and sufficient to reduce zygotic mating behaviors. Mei2 imposes the meiotic program and promotes the meiotic cycle, but also blocks mating behaviors independently of Mei3 and cell cycle progression. Thus, we find that fungi preserve zygote ploidy and survival by at least two mechanisms where the zygotic fate imposed by Mei2 and the cell cycle re-entry triggered by Mei3 synergize to prevent zygotic mating.

## Introduction

Most sexually reproducing species oscillate between haploid and diploid states. Meiotic divisions in diploid cells give raise to haploid progeny, while fertilization brings the genomes of two haploid partners together to form a diploid offspring. This ploidy cycle is central to the maintenance of a stable genome and relies on mechanisms that on the one hand promote faithful meiosis, and on the other hand ensure inheritance of exactly two parental genomes. Different species rely on processes that either restrict fusion to two parental gametes or two parental nuclei when fertilization occurs between multiple gametes (Beale et al., 2012; Snook et al., 2011; Wong and Wessel, 2005). Working on the fission yeast *Schizosaccharomyces pombe*, we recently discovered that fungal zygotes actively block recurrent fusion events. Upon fusion of the partner cells, the gamete-specific factors Mi and Pi form a bipartite transcription complex that triggers zygotic gene expression and blocks re-fertilization (Vještica et al., 2018) through mechanisms that remain unknown.

The fission yeast is a well-established model to probe the process of sexual differentiation. Mating is initiated when compatible partners from the two mating types, P and M, lack a source of nitrogen. Paracrine signaling between partners through diffusible pheromones and nitrogen depletion initiate the sexual development program dependent on the Ste11 transcription factor (Higuchi et al., 2002; Kunitomo et al., 2000; Sugimoto et al., 1991; Takeda et al., 1995). Increased Ste11 activity enhances expression of the pheromone and pheromone-signaling genes, which further drives gamete differentiation (Mata and Bähler, 2006). Starvation and pheromone signaling also promote arrest in the G1 phase of the cell cycle (Davey and Nielsen, 1994; Imai and Yamamoto, 1994). Several mechanisms concur in enforcing the G1 arrest, including 1) inhibition of CDK1-cyclin B complexes by cyclin-dependent kinase inhibitor, 2) cyclin degradation mediated by the anaphase promoting complex, and 3) reduction in cyclin mRNA stability (Navarro et al., 2017; Stern and Nurse, 1998, 1997). G1 arrest is essential for gamete differentiation, most likely as it is the only permissive phase for Ste11 to activate target genes (Kjaerulff et al., 2007; Shimada et al., 2008).

Once differentiated, gametes use the pheromone gradients released by the other mating type to search for a partner, grow towards each other, form pairs and fuse together (Bendezú and Martin, 2013; Merlini et al., 2016). In this process, the formin Fus1, whose transcription is induced by Ste11, is a major player, as it assembles the actin fusion focus at the site of cell-cell contact to drive local cell wall digestion and allow gametes to fuse together (Dudin et al., 2015; Petersen et al., 1995). Timely degradation of the cell wall precisely at the site of cell contact is critical since precautious perforations in the cell wall, which for instance occur upon pheromone signaling hyperactivation or upon construction of autocrine cells, lead to cell lysis (Dudin et al., 2016; Merlini et al., 2018). After gamete fusion, the fusion focus disappears, the fusion neck expands and the cells stop growing, the partner nuclei fuse together and the now diploid zygote immediately enters the meiotic cycle. Thus, even though the zygote is still in conditions of starvation and pheromone exposure, it fundamentally changes fate, reenters the cell cycle and blocks mating. This is all the more remarkable as pheromone signaling is essential for initiation of meiosis, at least in azygotic diploids (diploid cells not directly resulting from the fusion of two gametes; Tanaka et al., 1993).

The molecular details of meiosis initiation have been intensely studied in fission yeast (Figure 1A, Harigaya and Yamamoto, 2007). The principal target of the Mi-Pi bipartite transcription factor is the *mei3* gene, which is considered a master inducer of meiosis (McLeod et al., 1987; Vještica et al., 2018; Willer et al., 1995). Mei3, a small, largely disordered protein, functions as an inhibitor of the Pat1 kinase. Forced Mei3 expression, or direct inactivation of Pat1, are both sufficient to trigger meiosis even from a haploid genome (Cipak et al., 2014; McLeod and Beach, 1988). In haploids, which do not express *mei3*, Pat1 phosphorylates and inhibits the RNA-binding protein Mei2, whose activation is also sufficient to initiate meiosis (Harigaya and Yamamoto, 2007; Watanabe and Yamamoto, 1994). A large body of work, performed primarily on azygotic diploid cells using acute inactivation of Pat1 kinase, has dissected the critical role of Mei2 in driving meiosis. This showed that Mei2 is essential for both G1/S and, in association with the long non-coding *meiRNA*, G2/M meiotic transitions (Watanabe et al., 2001; Watanabe and Yamamoto, 1994). However, this work largely ignored the functions of Mei3, which was assumed to solely initiate a linear Mei3-Pat1-Mei2 pathway (Figure 1A). Consistent with this idea, indirect radioactive labelling of DNA suggested that *mei3Δ* zygotes become arrested before the initiation of premeiotic DNA replication (Egel and Egel-Mitani, 1974), as was reported for *mei2Δ* diploids (Watanabe and Yamamoto, 1994). We previously found that both *mei3Δ* and *mei2Δ* zygotes mate with a third gamete (Vještica et al., 2018). However, this phenotype is much more prominent in double mutants (Vještica et al., 2018), suggesting a non-linear pathway. These observations led us to investigate the roles of Mei3 and Mei2 in blocking re-fertilization and driving the meiotic cell cycle.

**Figure 1.**
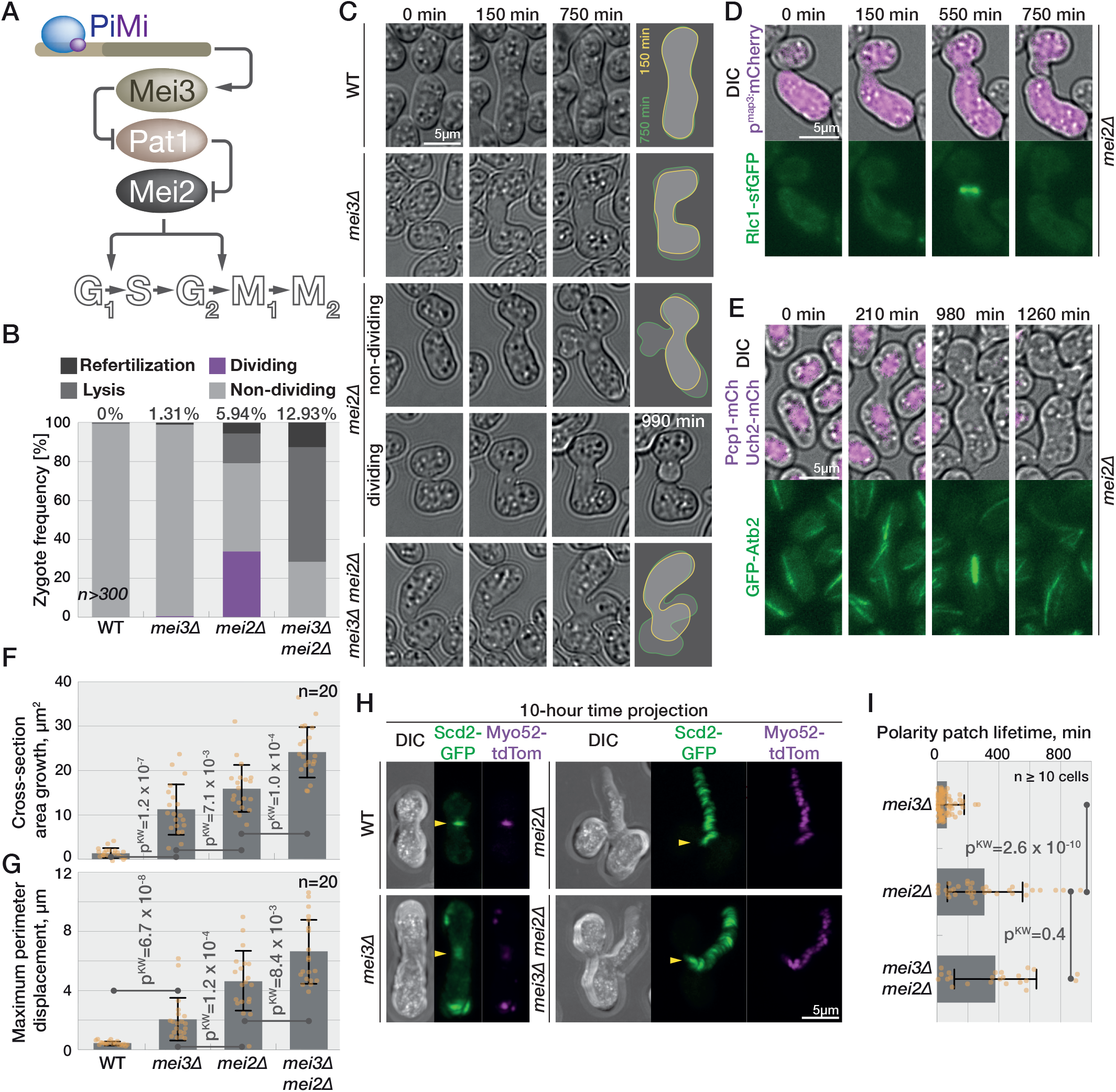
Meiotic signaling mutants show distinct phenotypes. **(A)** Schematic representation of meiotic signaling at the start of this study. **(B)** Quantification of phenotypes observed by long-term DIC imaging of zygotes. Numbers at the top report frequency of re-fertilization. **(C)** DIC images of cells prior to partner fusion (0 min), at the time zygotes complete concave neck expansion (150 min; yellow outline in right panel) and 10h later (750 min; green outline). A fraction of *mei2Δ* zygotes divides (here at 990 min). **(D)** Myosin light-chain Rlc1-sfGFP (bottom panel) in *mei2Δ* cells expressing mCherry from P-cell specific *p*^*map3*^ promoter (top panel, magenta overlaid with DIC) from pre-fusion (time 0) to zygote septation. **(E)** GFP-α-tubulin (bottom panel) in *mei2Δ* cells also expressing nuclear marker Uch2 and SPB marker Pcp1 fused to mCherry (top panel, magenta overlaid with DIC). Note the cell division after the zygote forms a microtubule spindle. **(F-G)** Area increase (F) and maximum perimeter displacement (G) of zygotes observed by time-lapse imaging during 10h after neck expansion measured from outlines as shown in (C). Dots show individual measurements, bars mean values and error bars standard deviation. p-values for indicated comparisons were obtained from the Krus-kal-Wallis test. **(H)** Projection of 60 timepoints during 10h from the time of neck expansion of polarity and growth markers Scd2-GFP (green) and Myo52-tdTomato (magenta). The arrowheads point to the signal at the neck. Both proteins are restricted to the neck in wildtype cells. In *mei2Δ* and *mei3Δ mei2Δ* zygotes both markers continuously trail the growth projection while *mei3Δ* zygotes show a fluorescent signal appearing at various cortical sites. **(I)** Quantification of Myo52-tdTomato patch lifetime from data as (H) (n ≥ 10 cells). Data is presented as in (F).

Here we show that zygotic mating blocks not only prevent re-fertilization but also avert lysis caused by autocrine signaling post-fertilization. Our findings reveal that zygotes block mating through two pathways, dependent on Mei3 and Mei2, respectively. First, we show that Mei3 promotes exit from the G1 phase, which by itself blocks re-fertilization. Second, we find that Mei2 confers the zygotic fate, which ensures not only meiotic commitment but also mating arrest, independently of cell cycle progression. Thus, fungal cells exhibit two pathways to block zygotic mating, ensuring the fidelity of the ploidy cycle.

## Results

### Mei3 and Mei2 independently regulate zygotic growth, cell division, lysis and mating

We previously reported that in addition to regulating meiotic entry, Mei3 and Mei2 also act to prevent zygotic mating and re-fertilization (Vještica et al., 2018). Indeed, deletion of either *mei3* or *mei2* led to re-fertilization in ~1% and ~6% zygotes, respectively (Figure 1B and Vještica et al., 2018). We also observed that simultaneous deletion of *mei3* and *mei2* genes increased the incidence of re-fertilizations to ~13% zygotes (Figure 1B and Vještica et al., 2018). These results suggested that Mei3 and Mei2, which initiate meiosis *via* a linear Mei3-Pat1-Mei2 cascade, may also act independently of each other to repress mating in zygotes.

We carefully analyzed the phenotypes of wildtype, *mei3Δ*, *mei2Δ* and *mei3Δ mei2Δ* zygotes (Figure 1C and Movie 1). After fertilization, wildtype zygotes expanded the fusion neck and underwent sporulation. The mutant zygotes also expanded the fusion necks but never sporulated and instead grew mating projections, re-fertilized and/or lysed. Surprisingly, we also found that one third of *mei2Δ* zygotes went through cell division, a phenotype that was virtually absent from *mei3Δ* and *mei3Δ mei2Δ* mutant zygotes (Figure 1B-C). Two lines of evidence suggest that these *mei2Δ* zygotes divide mitotically. First, dividing *mei2Δ* zygotes formed an actomyosin ring: after *mei2Δ* gametes fused, shown by the entry of cytosolic mCherry from the P-into the M-partner, the myosin light chain Rlc1-sfGFP formed a contractile ring in the zygotes, which subsequently underwent septation (Figure 1D and Movie 2). Second, dividing *mei2Δ* zygotes underwent nuclear division using a microtubule spindle, as shown by the nuclear marker Uch2-mCherry (Shaner et al., 2004) and α-tubulin GFP-Atb2 (Figure 1E). Spindle formation, nuclear division, contractile ring assembly and septation were not observed in *mei3Δ* and *mei3Δ mei2Δ* zygotes. We conclude that 1) in absence of Mei2 a fraction of zygotes progresses through mitosis, and 2) mitotic entry in *mei2Δ* zygotes depends on Mei3.

The only known molecular function of Mei3 is to inhibit the Pat1 kinase. We thus tested whether Mei3 signals through Pat1 to initiate mitotic entry in *mei2Δ* zygotes. Since *pat1* deletion is inviable due to constitutive Mei2 activation, we analyzed double and triple *mei3Δ*, *pat1Δ* and *mei2Δ* mutants (Figure S1A-B): ~37% *pat1Δ mei2Δ* double mutant zygotes underwent cell division, similar to *mei2Δ*. Again, zygotic cell division was virtually absent from *mei3Δ pat1Δ mei2Δ* zygotes. Thus, Mei3 promotes mitotic entry independently of Pat1 in *mei2Δ* zygotes.

Even when comparing zygotes that did not undergo mitosis, *mei3Δ* and *mei2Δ* single and double mutant zygotes showed clear differences in zygotic growth, morphology and lysis. To quantify zygotic growth and morphology we segmented cross-sections immediately after fusion necks have expanded (Figure 1C, 150 min timepoint and yellow line in the far right panel) and 10 hours later (Figure 1C, 750 min timepoint and green line in the far right panel). We then quantified the cross-section area increase and three morphological parameters of zygotic growth including 1) the maximum cell perimeter displacement, 2) the mean coefficient of variation in cell perimeter displacement, which is inversely linked to growth isometry, and 3) the diameter of the mating projections (Figures 1F-G, S1C-D). Growth was absent from wildtype zygotes between these two timepoints, whereas the cross-section area increased in all mutants (Figure 1F). The growth in *mei3Δ mei2Δ* zygotes was most prominent and clearly a consequence of cells forming long mating projections, which mostly originated from the fusion neck region (Figure 1C, 1F-G, S1C and Movie 1). Non-dividing *mei2Δ* zygotes exhibited a qualitatively similar behavior, but less extensive growth of mating projections. Conversely, *mei3Δ* zygotes did not form pronounced mating projections, and instead enlarged largely isometrically (Figure 1C, 1F-G, S1C-D and Movie 1). Deleting *pat1* in *mei2Δ* and *mei3Δ mei2Δ* did not change the overall phenotypes of these two genetic backgrounds (Figure S1B and Movie 3).

We further investigated the differences in growth and morphology of *mei2* and *mei3* mutant zygotes by monitoring the dynamics of the cell growth machinery. Specifically, we imaged Scd2-GFP, a scaffold protein that binds the active form of the key mating and growth regulator Cdc42 (Chang et al., 1994; Martin and Arkowitz, 2014). We simultaneously monitored Myo52-tdTomato, a type-V myosin that delivers secretory vesicles to growth and fusion sites (Bendezú and Martin, 2013, 2011; Dudin et al., 2015; Win et al., 2001). In wildtype cells, Scd2 and Myo52 focalized at the partner fusion site and then dispersed throughout the zygote after fertilization (Figure 1H and Movie 4). In *mei3Δ* zygotes, Scd2 and Myo52 signal also rapidly dispersed from the fusion site, but then formed relatively short-lived patches throughout the cell cortex (Figure 1H-I). The patches occasionally stabilized at the sites where mating projections formed (Movie 4). In *mei3Δ mei2Δ* and non-dividing *mei2Δ* zygotes, Scd2 and Myo52 patches were significantly more stable after fertilization and marked the growing mating projection (Figure 1H-I and Movie 4). In these zygotes, Scd2 and Myo52 patches frequently did not disassemble after fertilization and mating projections grew from the region of the fusion neck. These differences in zygotic growth between *mei2Δ* and *mei3Δ* support the conclusion that Mei2 has roles in regulating the zygotic growth machinery that are independent of upstream Mei3 signaling.

Taken together, our analyses of individual and double *mei2Δ* and *mei3Δ* mutants show that, in addition to their well-established co-dependent role in meiotic induction, Mei2 and Mei3 independently regulate the zygotic cell cycle, growth and mating blocks: 1) Mei3 promotes cell cycle progression in absence of Mei2, 2) Mei2 prevents zygote mating and lysis even in absence Mei3, and 3) mating and lysis phenotypes are exacerbated in the double mutant.

### Mating blocks prevent zygote lysis

Lysis is a prominent phenotype of *mei3Δ mei2Δ* zygotes that we also readily observed in *mei2Δ* zygotes but not in mutant gametes prior to fertilization (Figure S2A). What causes lysis specifically post-fertilization? Cell wall digestion is normally brought about by paracrine pheromone signaling and formation of an actin fusion focus nucleated by the formin Fus1 at the site of partner contact (Dudin et al., 2015; Martin, 2019). In self-stimulating autocrine cells, high levels of pheromone signaling induce the formation of an ectopic fusion focus, which leads to localized cell wall degradation independently of a partner cell, and consequently lysis (Dudin et al., 2016). Lysis in autocrine cells is suppressed by deletion of the formin Fus1 (Dudin et al., 2016). As gamete fusion brings together the machineries for production, secretion and perception of both pheromones, we hypothesized that zygotes behave as autocrine cells, which would explain their frequent lysis.

To test whether the death of *mei3Δ mei2Δ* zygotes is a consequence of failed fusion attempts, we aimed to deplete Fus1 specifically post-fertilization by developing a zygote-specific artificial degron system (Figure 2A). We first generated DegGreen by fusing the N-terminus of the ubiquitin E3 ligase complex protein Pof1 with the GFP-binding protein (GBP) and expressing it from a strong constitutive promoter. We reasoned that GFP-GBP binding will direct the Pof1 E3 ligase activity towards GFP and lead to its degradation. Indeed, introducing the DegGreen into cells expressing meGFP (monomeric enhanced GFP) lowered the green fluorescent signal by 95% (Figure 2B). We then built DegRed, a fusion between Pof1 N-terminus and mCherry-binding protein, which decreased the red fluorescence in cells expressing mCherry by 63% (Figure 2C). When we introduced DegGreen into gametes that carried the native Fus1 fused with meGFP, the fusion efficiency to *fus1Δ* partners was reduced by ~5-fold as compared to control cells carrying individual proteins (Figure 2D). Likewise, expression of DegRed in cells with native Fus1 fused to mCherry reduced fusion efficiency to *fus1Δ* partners by ~2-fold as compared to cells carrying individual proteins (Figure 2E). Thus, both artificial degrons can decrease the function of Fus1 in gametes.

**Figure 2.**
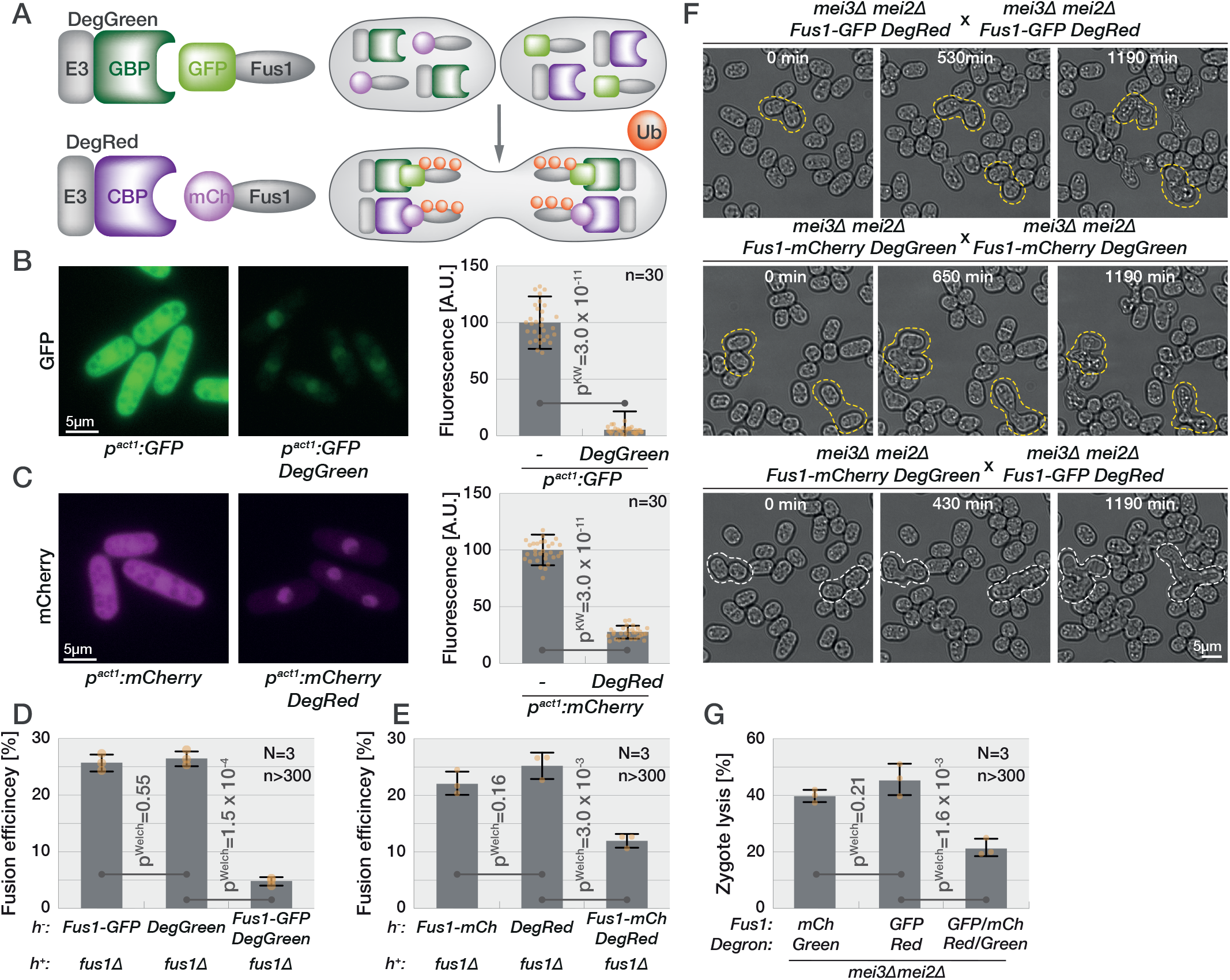
Zygote lysis is induced by aberrant fusion attempts. **(A)** Schematic of the artificial degron system. The DegGreen construct composed of the N-terminal, E3 ligase domain of Pof1 (E3, grey) fused with the GFP-binding nanobody (GBP, dark green) targets GFP (light green; here linked to Fus1) for degradation. Similarly, the DegRed construct, consisting of the E3 ligase domain of Pof1 (E3, grey) fused with the mCherry-binding nanobody (CBP, dark purple) targets mCherry (light purple) for degradation. For zygote-specific protein depletion, one partner gamete carries DegGreen and the mCherry-tagged protein of interest, while the other partner brings DegRed and the GFP-tagged protein of interest. Ubiquitination (orange circles) and degradation only happens post-fertilization. **(B-C)** Exponentially growing cells expressing GFP (B) or mCherry (C) from the strong *p*^*act1*^ promoter without (left) or with DegGreen (B) or DegRed(C) (right). The graph shows the fluorescence normalized to cells without degron systems. Dots show individual measurements, bars mean values and error bars standard deviation. p-values for indicated comparisons were obtained from the Kruskal-Wallis test. **(D-E)** Gamete fusion efficiency between *h-* strains with incomplete or complete (right) Fus1 degron systems and *h+ fus1Δ.* Dots show individual replicates, bars mean values and error bars standard deviation. p-values for indicated comparisons were obtained from the Welch’s test. **(F)** Mating of *mei3Δ mei2Δ* gametes carrying the indicated halves of the Fus1 artificial degron system. Zygotes with the incomplete Fus1 degron (top and middle panels) undergo cell lysis (examples outlined in yellow), whereas zygotes with a complete Fus1 degron post-fertilization (bottom panel) prevent lysis (examples outlined in white). **(G)** Frequency of lysis in *mei3Δ mei2Δ* zygotes with incomplete or complete degron systems (genotypes as in (F). Data is presented as in (D).

To specifically degrade Fus1 in zygotes, we used the *mei3Δ mei2Δ* genetic background and introduced DegGreen into gametes expressing Fus1-mCherry, and DegRed into the other gamete expressing Fus1-meGFP, such that substrates and degrons come in contact only post-fertilization (Figure 2F and Movie 5). Live imaging showed high levels of either Fus1-GFP or Fus1-mCherry in control *mei3Δ mei2Δ* zygotes lacking the cognate artificial degron, with fluorescence concentrated at the shmoo tip, similar to the Scd2 and Myo52 localization described above (Figure S2B, top and middle panels).

Remarkably, zygotes with both degrons and Fus1 tags showed reduced Fus1 fluorescence post-fertilization and decreased cell lysis by at least 2-fold relative to control zygotes (Figure 2G, S2C). Thus, cell death in *mei3Δ mei2Δ* mutant zygotes is a consequence of their inability to block mating. We conclude that mating blocks prevent zygote death.

### Mei3 promotes G1-S cell cycle transition independently of Pat1 and Mei2

Our findings that *mei2Δ*, but not *mei3Δ mei2Δ*, zygotes undergo mitosis strongly suggests that Mei3 drives cell cycle progression post-fertilization. To directly assess progress through the cell cycle, we used the Hoechst 3334 dye to stain the DNA of cell mixtures produced by mating of heterothallic *h+* (P) and *h-* (M) strains carrying either cytosolic sfGFP or mCherry expressed from the strong *p*^*tdh1*^ promoter (Figure 3A). The two fluorophores distinguish gametes, which fluoresce in a single channel, from zygotes, which fluoresce in both channels using flow cytometry, and enabled us to determine the cell cycle stage of each cell type in mating mixtures. We performed this experiment in *mei3Δ*, *mei2Δ*, *mei3Δ mei2Δ*, and as control *mei4Δ*, which arrest with 4C DNA content prior to sporulation (Horie et al., 1998; Murakami-Tonami et al., 2007). As expected, all mutant gametes arrested with 1C content in response to nitrogen starvation and mating (Figure 3A, magenta and green curves). Mutant zygotes however exhibited differences in DNA content (Figure 3A, yellow curves). Consistent with previous work, *mei4Δ* zygotes arrested with 4C content. Approximately half of the *mei2Δ* zygotes also had 4C DNA content, indicating genome replication. The exact number of *mei2Δ* zygotes that progress to S-phase could not be determined in this manner since some zygotes pass mitosis prior to our flow cytometry measurements (Figure 1B). By contrast, *mei3Δ* and *mei3Δ mei2Δ* zygotes arrested with 2C content, indicating that fertilization brought partner genomes together but the zygotes did not replicate their genomes. We obtained similar results in experiments with homothallic (*h90*; self-fertile) strains where mCherry expression was driven by the P-cell specific *p*^*map3*^ promoter, and sfGFP by the M-cell specific p^mam1^ promoter (Figure S3A). Zygotes produced by *h90 mei3Δ* and *mei3Δ mei2Δ* strains arrested the cell cycle in G1 phase, while a large fraction of *mei2Δ* zygotes progressed through S-phase. In this experiment, we used as control *sme2Δ* mutants, which lack the Mei2-binding *meiRNA* and arrest with 4C DNA content prior to sporulation (Watanabe and Yamamoto, 1994). We conclude that Mei3 promotes S-phase entry in zygotes.

**Figure 3.**
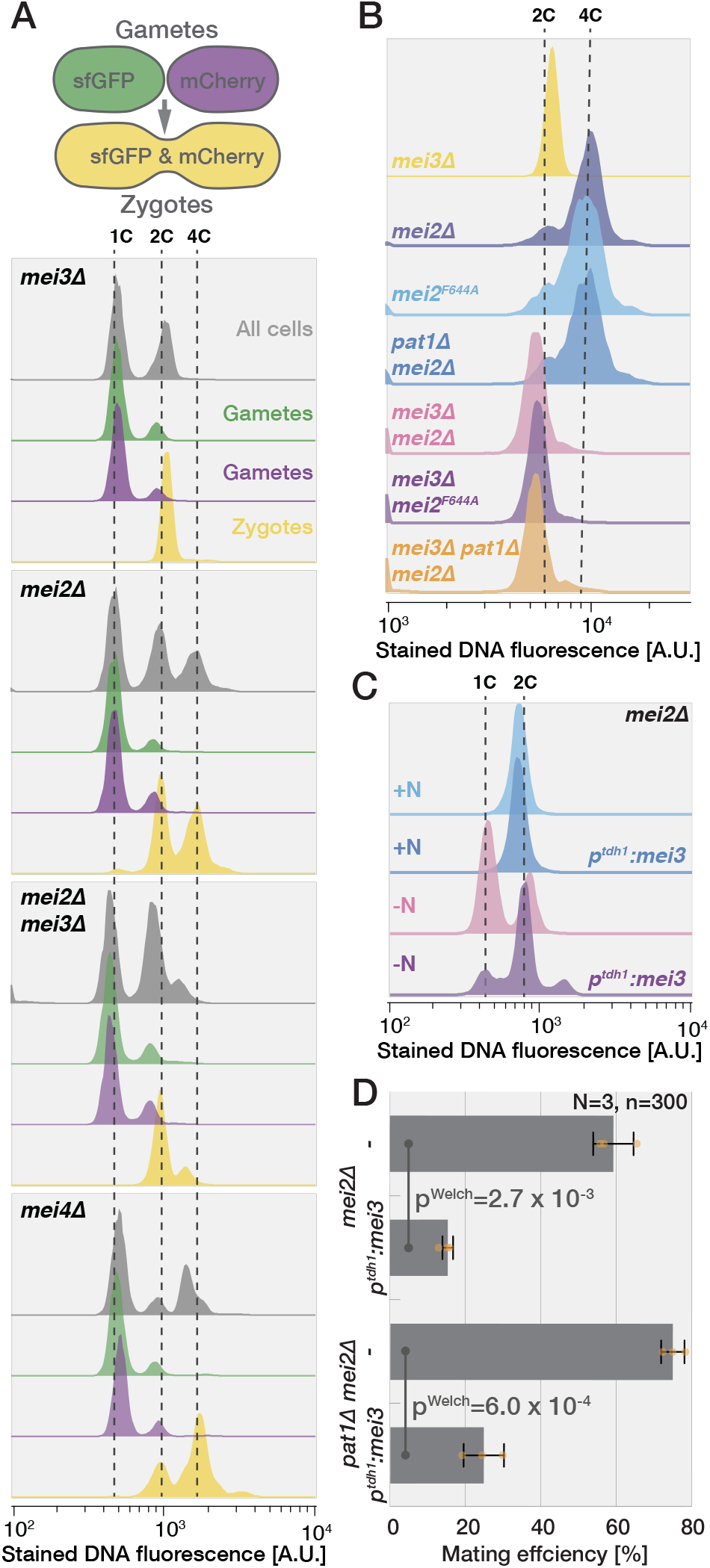
Mei3 promotes G1 exit independently of Pat1 and Mei2. **(A)** Strategy used for flow cytometry on mating mixtures of heterothallic strains that constitutively express either sfGFP or mCherry (top). Gametes, which carry a single fluorophore, can be distinguished from zygotes, which carry both fluorophores. Flow cytometry analysis of Hoechst-stained DNA fluorescence (x-axis) in mating mixtures after 24h (bottom). The y-axis shows the cell number normalized to mode. Profiles of all cells (grey), gametes (green and magenta) and zygotes (yellow) are shown. Gametes largely arrest with unreplicated 1C genomes. *mei3Δ* and *mei3Δmei2Δ* zygotes show unreplicated 2C content. *mei2Δ* and *mei4Δ* zygotes show a prominent 4C peak. **(B)** Hoechst-stained DNA fluorescence in zygotes identified through gating for increased size in forward and side scatter measurements (see Materials and Methods). Zygotes lacking *mei3* arrest with unreplicated genomes (2C), while those with intact *mei3* replicate their genomes even in absence of *mei2* and *pat1*. **(C)** Hoechst-stained DNA fluorescence of haploid *mei2Δ* cells with or without *mei3* expressed from the constitutive *p*^*tdh1*^ promoter grown in presence or absence of nitrogen for 24h. Ectopic expression of Mei3 prevents the G_1_-arrest in nitrogen starved gametes but has no effect in nitrogen-rich media. **(D)** Mating efficiency of *mei2Δ* and *mei2Δ pat1Δ* cells with or without *mei3* expressed from the constitutive *p*^*tdh1*^ promoter after 24h incubation on agar plates. Constitutive *mei3* expression reduces mating of gametes independently of *mei2* and *pat1*. Dots show individual replicates, bars mean values and error bars standard deviation. p-values for indicated comparisons were obtained from the Welch’s test.

We further tested whether Mei3 function in driving the S-phase entry relies on Pat1. To that aim we performed flow cytometry measurements of DNA content in zygotes produced by mutants lacking individual and combinations of *mei3, mei2 and pat1* genes. We note that in this instance we did not label the gametes with fluorescent proteins since we found that we could use the flow cytometry forward and side scatters to separate zygotes based on their larger size (Figure S3B, see Materials and Methods). Removal of Pat1 had no effect on the ability of zygotes to enter S-phase as both *mei2Δ* and *mei2Δ pat1Δ* zygotes accumulated 4C content while all zygotes deleted for *mei3* arrested in G1-phase (Figure 3B). Mei3-driven DNA replication also occurred in zygotes where Mei2 RNA-binding was abolished due to the F644A mutation in the RRM motif (Figure 3B, Watanabe and Yamamoto, 1994). We conclude that Mei3 drives G1-S transition in zygotes independently of Mei2 and Pat1.

Though independent of downstream zygotic signaling, Mei3 may require some zygote-specific factors to promote S-phase entry. To test this hypothesis, we ectopically expressed Mei3 from the strong *p*^*tdh1*^ promoter in haploid cells that remained alive due to simultaneous deletion of the *mei2* gene. During exponential growth in nitrogen rich media, overexpression of Mei3 did not cause detectable defects in DNA replication based on flow cytometry measurements (Figure 3C). However, upon nitrogen removal, which arrested the *mei2Δ* haploids in G1-phase, a large fraction of cells overexpressing Mei3 had duplicated genomes. In agreement with impaired ability to arrest in G1, overexpression of Mei3 resulted in decreased mating efficiency (Figure 3D). Mei3 overexpression impaired mating also in cells lacking Pat1 consistent with our results that Mei3 cell cycle effects are Pat1-independent (Figure 3D). We conclude that Mei3 functions independently of zygote-specific factors to promote the G1-S transition.

### Mei3 promotes S-phase entry and represses mating in diploid fission yeast

Our results contrast earlier work that proposed that Mei2 is required for S-phase entry in nitrogen-starved diploid cells committed to azygotic meiotic differentiation (Harigaya and Yamamoto, 2007; Watanabe and Yamamoto, 1994). Note that the diploids in these experiments are to be distinguished from zygotes as they do not directly result from the fusion of two haploid cells but have been propagated mitotically as diploids before meiotic induction by nitrogen starvation. Diploids are typically isolated from mating mixtures of two haploid strains bearing different *ade6* mutant alleles that exhibit intermolecular complementation (Ekwall and Thon, 2017). We decided to revisit these findings and prepared wildtype and diploid cells lacking *mei4* or, either individually or combined, *mei2* and *mei3* (Figure S4A). Since pheromone response is necessary for Mei3 induction and downstream meiotic differentiation (McLeod et al., 1987; Tanaka et al., 1993; Willer et al., 1995), we also monitored the expression of mCherry and sfGFP placed under the pheromone-responsive, mating type-specific *p*^*map3*^ and *p*^*mam1*^ promoters, respectively (Figure S4A). Importantly, we observed that even the wildtype diploid population is heterogeneous with 1) cells that do not induce expression from both *p*^*map3*^ and *p*^*mam1*^ (Figure S4A, outlined cell), 2) cells that mate (Figure S4A, arrow) and 3) cells that respond to nitrogen removal asynchronously. These issues result in flow cytometry subpopulations (Figure S4C) and complicate the analysis of flow cytometry data.

To overcome these issues, we used two complementary approaches. First, we adapted the flow cytometry gating to evaluate the DNA content only in small cells, which excluded mated and clumped cells (see Materials and Methods). We also restrained the analysis of nitrogen starved cells to those with clear *p*^*map3*^ and *p*^*mam1*^ promoter activity required for meiotic differentiation (Figure S4B). Before nitrogen starvation, a majority of cells had replicated genomes. Whereas wildtype, *mei2Δ* and *mei4Δ* cells showed clear G1 and G2 cell populations after 24h of nitrogen starvation, the *mei3Δ* and *mei3Δ mei2Δ* diploids completely arrested in G1 phase within 12 hours after nitrogen removal. In a second approach, we argued that *mat1-P* and *mat1-M* loci are not stable in diploids cell produced by standard *h+* and *h-* strains (Beach and Klar, 1984), a hypothesis in agreement with our observation of diploid cells with a single active mating type-specific promoter (Figure S4A, outlined cell). To overcome this problem, we produced diploids where *mat1* locus switching was prevented by a *H1Δ17* mutation (Arcangioli and Klar, 1991; Vještica et al., 2018). These diploids showed activity from both *p*^*map3*^ and *p*^*mam1*^ promoters, did not mate, and sporulated within 24 hours after nitrogen removal (Figure 4A). Flow cytometry analysis also showed increased homogeneity and similar activation of both *p*^*map3*^ and *p*^*mam1*^ promoters (Figure S4C, note that cells largely fall along the diagonal). Flow cytometry measurements of DNA content on all non-clumped *mat1-H1Δ17* diploids (70%-95% of all cells) clearly showed that *mei3Δ* and *mei3Δ mei2Δ* diploids completely arrested in G1, while wildtype, *mei2Δ* and *mei4Δ* cells replicated their genomes (Figure 4B). Furthermore, imaging revealed that ~10% *mei2Δ* diploids, but no other genotype, underwent cell division after cells had activated the two pheromone responsive promoters (Figure 4C, 4F, Movie 6). Thus, nitrogen-starved azygotic diploids, like zygotes, require Mei3 for the G1-S transition.

**Figure 4.**
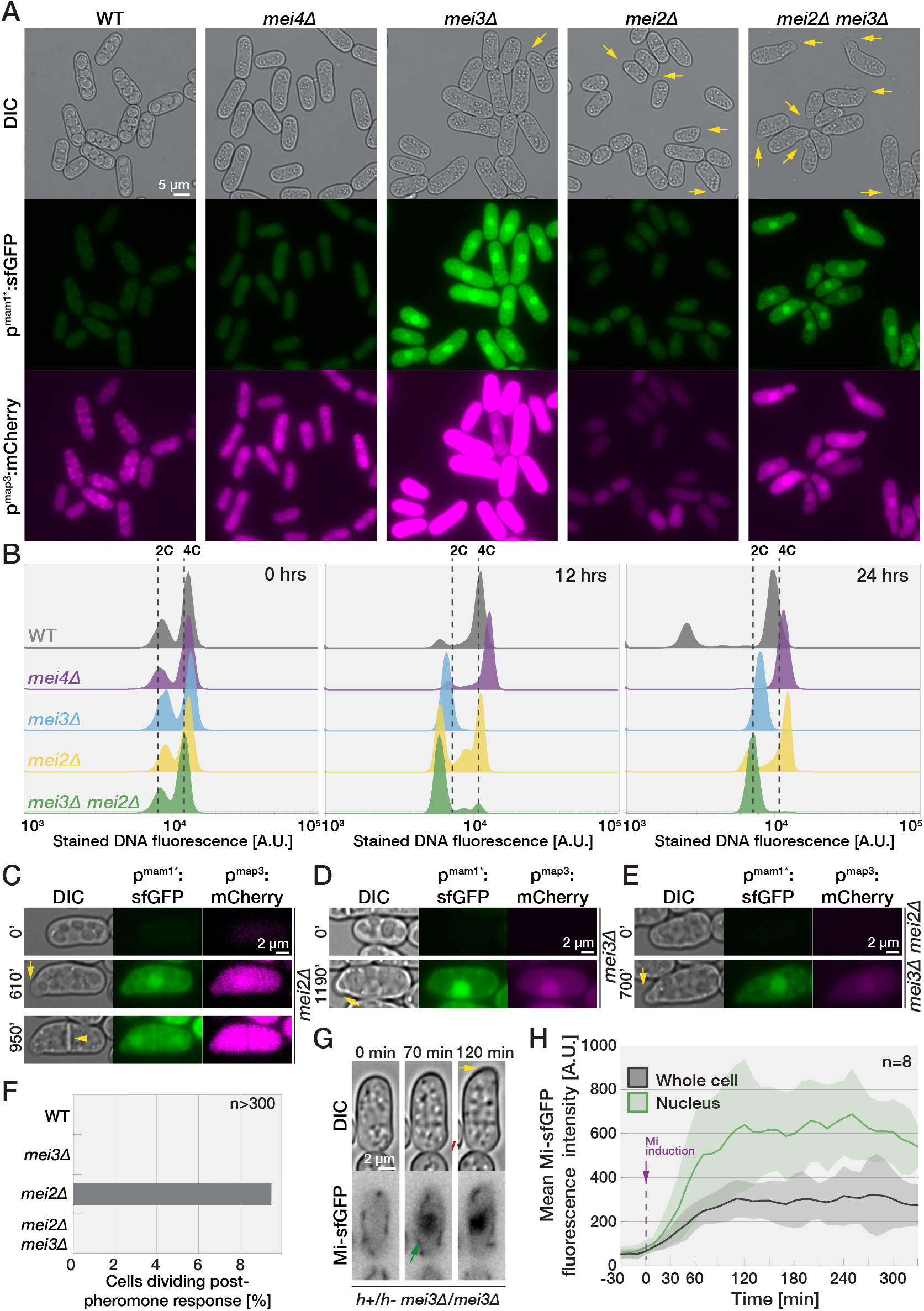
Mei3 promotes G1 exit in stable diploid cells. **(A)***H1Δ17* diploid cells expressing mCherry and sfGFP from P- and M-cell specific promoters *p*^*map3*^ and *p*^*mam1*^ 24h after removal of nitrogen. Arrows point to shmoo-like projections in *mei3Δ* and *mei2Δ* mutants. **(B)** Flow cytometry analysis of Hoechst-stained DNA fluorescence (x-axis) in diploid cell used in (A) at indicated timepoints after nitrogen removal. The y-axis shows the cell number normalized to mode. Note that *mei3Δ* and *mei3Δ mei2Δ* mutants arrest with unreplicated genomes (2C) while wildtype, *mei4Δ* and *mei2Δ* cells replicate their DNA (4C). The sub-2C population in wildtype sample are likely spores. **(C-E)** Time-lapses show induction of fluorophores in cells as in (A). Note formation of mating projections (arrows) and the septum in *mei2Δ* mutant (arrowhead). **(F)** Percentage of *H1Δ17* diploids expressing both *p*^*mam1*^:GFP and *p*^*map3*^:mCherry that are septating. **(G-H)** Time-lapse (G) and quantification (H) of Mi-sfGFP induction in *H1Δ17* diploid cells deleted for *mei3*. Note that Mi-sfGFP induction (black line in H) coincides with its nuclear accumulation (green arrow in G and green line in H) and precedes the formation of the shmoo (yellow arrow in G). The graph reports mean fluorescence for 8 cells with shaded areas denoting standard deviation.

While nitrogen starved otherwise wildtype and *mei4Δ mat1-H1Δ17* diploids did not grow, *mei2Δ* and/or *mei3Δ* diploids formed shmoo-like projections (Figure 4A, arrows). Time-lapse microscopy revealed that mating projections in *mei2Δ*, *mei3Δ* and double mutant diploids formed after activating both P-and M-pheromone-responsive promoters (Figure 4C-E and Movie 6). These results imply that, like in zygotes, Mei2/Mei3 signaling blocks mating in diploid cells starved for nitrogen.

These observations raise a conundrum about the role of pheromone signaling in diploids. Indeed, expression of the Pi-Mi complex essential for Mei3 and thus meiotic induction relies on pheromone signaling (Willer et al., 1995), yet our results show that Mei3 actively represses pheromone-induced mating in diploids. We reasoned that induction of the Pi-Mi complex may occur at lower levels of pheromone signaling than needed to form mating projection. Indeed, expression of Mi-sfGFP in *mei3Δ* mutant diploids was observed prior to formation of mating projections (n>50 diploids forming a shmoo, Figure 4G and Movie 7). Appearance of the Mi-sfGFP signal though asynchronous in the population was concomitant with its nuclear accumulation (Figure 4G-H), which depends on Pi (Vještica et al., 2018 and Figure S4D-E). This suggests that the Pi-Mi complex forms at the time of Mi induction. These results are consistent with the view that in diploids, low levels of pheromone signaling induce the Pi-Mi complex, leading to Mei3 expression, which in turn represses further pheromone-dependent mating and initiates meiosis.

We conclude that in diploid cells, like in zygotes, Mei3 promotes the G1-S transition and both Mei2 and Mei3 repress mating behaviors.

### Mei3 promotes the premeiotic S-phase

We showed above that Mei3 can advance the cell cycle in mitotic cells and is required for premeiotic S-phase in diploids and zygotes. However, interpreting the function of Mei3 in premeiotic S-phase is complicated by the well-established role of Mei3 in relieving the Pat1-mediated inhibition of Mei2. To uncouple possible functions of Mei3 in promoting premeiotic S-phase from Mei2 activation, we built cells with a simplified <u>m</u>eiotic <u>s</u>ignaling that we refer to as *SMS* cells (Figure 5A). We reasoned that *pat1* deletion would be made viable in haploid cells by deletion of *mei2*, and that Mei2 expression can be induced specifically post-fertilization to induce meiosis. Accordingly, we deleted the *mei2* and *pat1* loci and replaced the native *mei3* open reading frame (which is induced post-fertilization) with that of *mei2*. *SMS* cells did not exhibit any obvious defects during exponential growth (data not shown) and produced zygotes that sporulated at wildtype rates (Figure 5B-C; p^Welch’s test^=0.28). We note that ~3±2% *SMS* zygotes produced asci with aberrant spore number, and that ~2±1% *SMS* zygotes died prior to sporulation. We also compared the levels of meiotic recombination between wildtype and *SMS* zygotes (Figure 5D). Briefly, strains carrying constructs for expression of cytosolic sfGFP or mCherry integrated at the *leu1* and *aha1* genomic loci, respectively, were mated and the proportion of progeny carrying each fluorophore was quantified. The genetic distance between the two markers decreased from 17.13±0.14 cM in wildtype to 14.70±0.13 cM in *SMS* mutants. We conclude that *SMS* zygotes, which lack Mei3, show minor overall defects, but no gross perturbation of the meiotic cycle.

**Figure 5.**
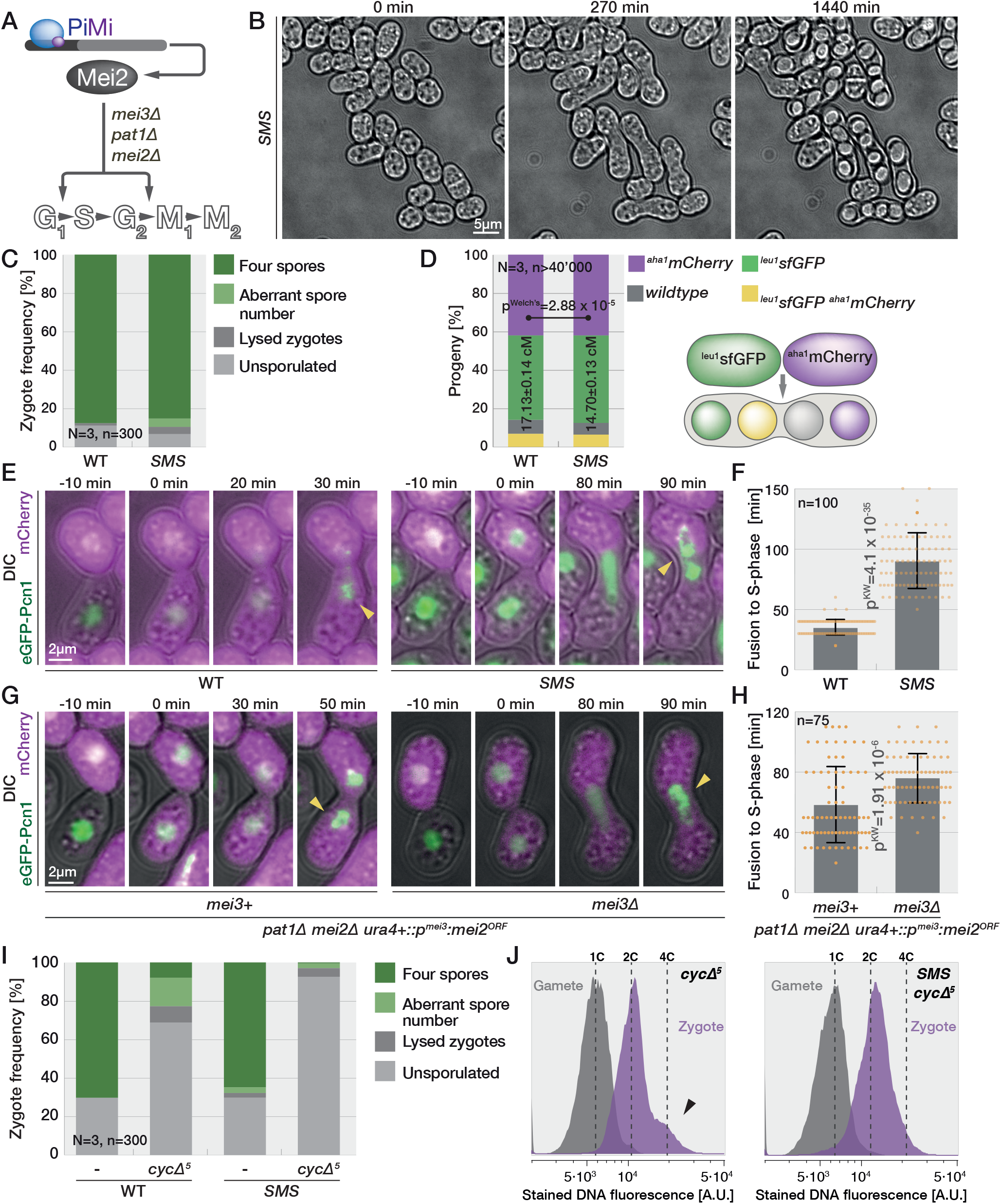
Mei3 advances premeiotic-S phase entry. **(A)** Schematic representation of meiotic signaling in *SMS* zygotes. **(B)** Time-lapse of *SMS* cells mating. **(C)** Quantification of zygotic phenotypes after 48h on solid media lacking nitrogen. **(D)** Results of flow cytometry quantification for non-recombinant (purple and green) and recombinant (grey and yellow) progeny obtained by crossing strains coding for sfGFP at the *leu1* locus with strains coding for mCherry at the linked *aha1* locus. **(E, G)** Time-lapses show DIC (grey) and GFP-Pcn1 (green) during mating of cells with indicated genotypes. The fusion time (0 min) is evident as partner exchange cytosolic mCherry (magenta) expressed in only one gamete. Punctate GFP-Pcn1 signal (arrowheads) is indicative of S-phase. Note that SMS zygotes take longer to exit G1 than wildtype (E), and that Mei3 promotes S-phase entry in zygotes lacking Pat1 (G). **(F,H)** Quantification of post-fertilization time to appearance of GFP-Pcn1 puncta determined from time-lapses as in (E) and (G). Dots show individual measurements, bars mean values and error bars standard deviation. p-values for indicated comparisons were obtained from the Kruskal-Wallis test. **(I)** Quantification of zygotic phenotypes after 24h on solid media lacking nitrogen. **(J)** Flow cytometry analysis of Hoechst-stained DNA fluorescence (x-axis) for gametes (grey) and zygotes (magenta) in mating mixtures after 18 h. The y-axis shows the cell number normalized to mode. Gametes arrest with unreplicated 1C genomes. Genome duplication occurs in a fraction of *cycΔ5* (left panel, arrow points a 4C shoulder) but not *SMS cycΔ5* zygotes (right panel).

*SMS* zygotes, however, showed a substantial delay in G1-S transition. We imaged wildtype and SMS zygotes carrying GFP fused to the N-terminus of the DNA replication fork component Pcn1 (Meister et al., 2007) expressed in addition to the native protein. Outside S-phase the GFP-Pcn1 produces a uniform nuclear signal but forms prominent foci during DNA replication (Meister et al., 2007; Vještica et al., 2020). Strikingly, GFP-Pcn1 foci formed before nuclear fusion in wildtype zygotes, but only after karyogamy in *SMS* zygotes (Movie 8). The time between partner fusion (monitored by entry of cytosolic mCherry in the P-partner) and S-phase onset was 91±23 min in *SMS* zygotes, but only 35±7min in wildtype zygotes (Figure 5E-F). To test whether Mei3 advances S-phase in cells with contracted zygotic signaling, we compared *mei3+* and *mei3Δ* cells, which had *pat1* and *mei2* loci deleted and *mei2* coding sequence under *p*^*mei3*^ promoter control at the *ura4* locus. Mei3 expression reduced the time between fertilization (marked here by entry of cytosolic mCherry in the M-partner) and premeiotic S-phase entry, labelled by Pcn1 foci, by ~17 minutes (Figure 5G, 5H). Thus, Mei3 promotes but is not essential for the premeiotic S-phase in zygotes with functional Mei2. This function is also independent of Pat1 kinase.

In wildtype zygotes, previous work showed that the meiotic G1-S transition is largely driven by additive effects of the five non-essential cyclins, Cig1, Cig2, Puc1, Crs1 and Rem1 (Gutiérrez-Escribano and Nurse, 2015). However, even upon deletion of these five cyclins (named here *cycΔ5*), ~23% zygotes sporulate (Figure 5I and Gutiérrez-Escribano and Nurse, 2015). Examination of DNA content in *cycΔ5* zygotes by flow cytometry showed a 2C peak with a clear 4C shoulder, indicating that a significant number of zygotes have sufficient residual CDK1 activity for premeiotic S-phase, likely driven by the remaining, essential B-type cyclin Cdc13 (Figure 5J). By contrast, premeiotic S-phase was almost completely blocked when combining the *cycΔ5* with the SMS background, which lacks Mei3, as shown by absence of the 4C shoulder in flow cytometry analysis, and only 3±0.5% zygotes sporulated (Figure 5I-J). The additive effects between *cycΔ5* and *SMS* genetic backgrounds further support the view that Mei3 serves to elevate cyclin-CDK activity to promote S-phase entry.

### Mei3-driven cell cycle progression prevents zygote mating and re-fertilization

Because Mei3 shows Mei2-independent roles in both promoting S-phase entry and blocking re-fertilization, we tested whether cell cycle progression *per se* may block re-fertilization. This hypothesis is also suggested from the knowledge that haploid cells mate only during the G1-phase of the cell cycle. To address this possibility, we monitored the timing of shmoo formation relative to S-phase entry in mating mixtures of zygotic signaling mutants. Time-lapse imaging of GFP-Pcn1 faithfully reproduced our earlier flow-cytometry findings with Pcn1 replication foci forming in virtually all wildtype, *sme2Δ* and *mei4Δ* zygotes but in none of the *mei3Δ* or *mei3Δ mei2Δ* zygotes (Figure S5A and Movie 9). In *mei2Δ* zygotes, we observed 50% passing S-phase, 37% arresting in G1-phase for the duration of the experiment and the remaining 13% lysing. Remarkably, all lysing cells were in the G1 phase, and the remaining G1-arrested *mei2Δ* zygotes formed mating projections (Figures 6A-B and Movie 10). Conversely, only 40% of zygotes that advanced the cell cycle formed shmoos and did so prior or during S-phase but never after S-phase completion. Thus, cell cycle progression correlates with mating repression in zygotes.

**Figure 6.**
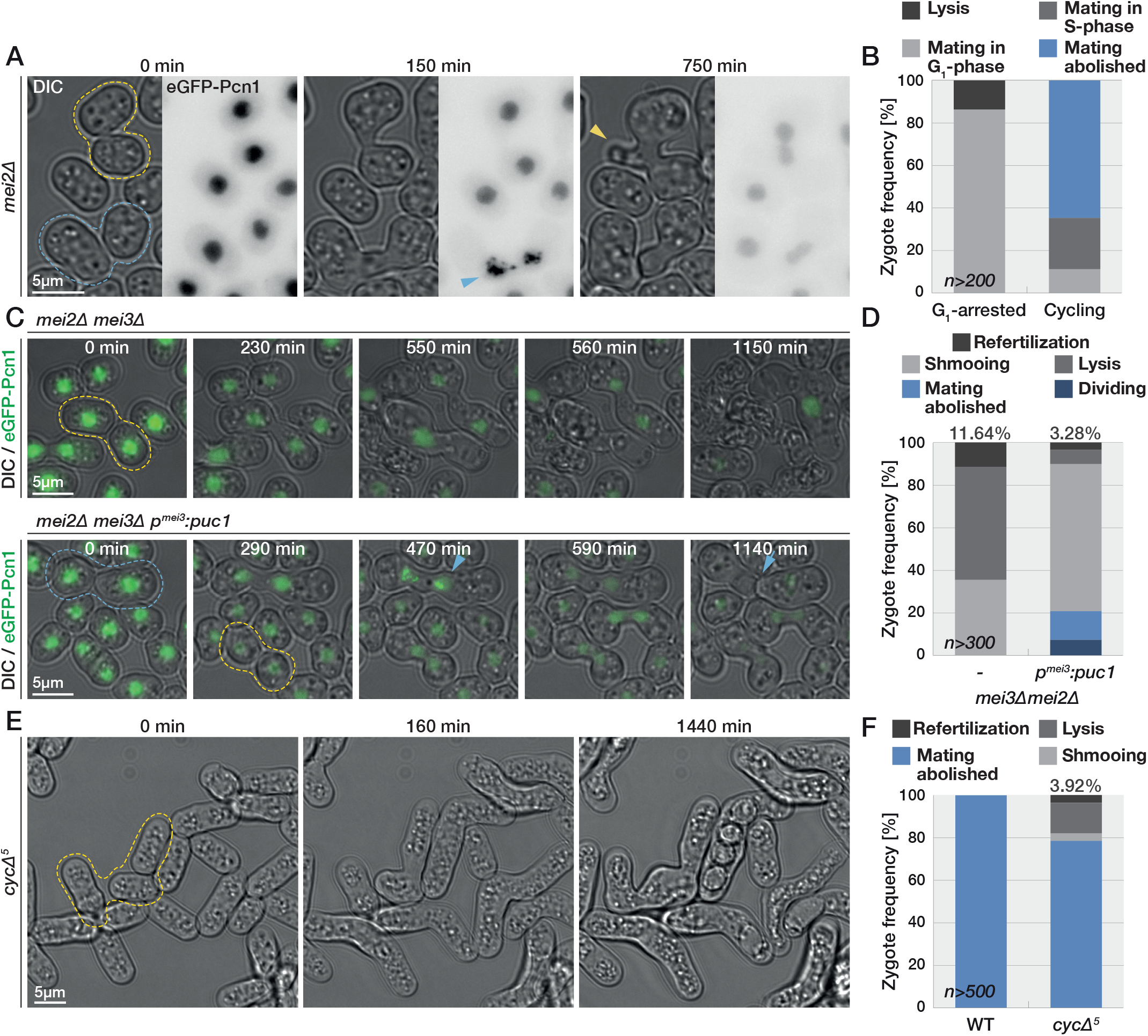
Cell cycle progression prevents re-fertilization. **(A)** Time-lapse of mating *mei2Δ* cells. Zygotes that persist in G1 (yellow outlines) form prominent mating projections (yellow arrowhead). Zygotes that progress through S-phase (blue outlines), evident from GFP-Pcn1 puncta (blue arrowhead), largely repress shmoo formation and instead divide. **(B)** Quantification of phenotypes of *mei2*Δ zygotes presented in (A). We used GFP-Pcn1 to group zygotes into those that either arrest (left bars) or exit (right bars) G1-phase and scored the mating behaviours of both groups. **(C)** Mating of control *mei3Δ mei2Δ* cells (top panel) and *mei3Δ mei2Δ* cells that express Puc1 from the zygote specific p^mei3^ promoter. GFP-Pcn1 (green) forms puncta (blue arrowhead) in Puc1-expressing zygotes (blue outlines). G1-arrested zygotes (yellow outlines) frequently lyse in control but not Puc1-expressing zygotes. **(D)** Quantification of zygotic phenotypes observed in time-lapses as in (D). **(E)** Mating of *cycΔ5* cells, which undergo re-fertilization (yellow outlines). **(F)** Quantification zygotic phenotypes observed in (E).

To test whether forced S-phase entry can suppress zygotic mating independently of Mei3 and Mei2, we used *mei3Δ mei2Δ* cells, which produce G1-arrested zygotes (Figure 3A, S5A), and forced the expression of the G1 cyclin Puc1 under the regulation of the zygote-specific *p*^*mei3*^ promoter. Imaging of the GFP-Pcn1 showed that forced expression of Puc1 postfertilization drove ~15% of zygotes through S-phase (Figures 6C, S5B and Movie 11) and ~7% through division (Figure 6D). Consistent with the hypothesis that cell cycle progression acts as a re-fertilization block, Puc1 expression decreased the rates of re-fertilization and lysis in *mei3Δ mei2Δ* mutant zygotes (Figures 6C-D, S5C). Puc1 expression also diminished the mating-induced growth in *mei3Δ mei2Δ* zygotes that did not initiate S-phase (Figure S5C). Our results imply that increased CDK activity and associated cell cycle progression is sufficient to prevent mating and re-fertilization in zygotes.

We next sought to test whether the G1-exit is necessary to prevent zygotic mating. To block otherwise wildtype zygotes in G1, we first used the analogue-sensitive CDK allele Cdc2-asM17, whose kinase activity can be inhibited by the ATP-analogue 1NM-PP1 (Aoi et al., 2014). After starving *cdc2-asM17* P- and M-gametes, expressing either cytosolic sfGFP or mCherry, separately to arrest cells in G1-phase, we applied high levels of the inhibitor 1NM-PP1 (25 μM), or DMSO as control, and mixed the gametes to allow mating. DMSO-treated zygotes went on to sporulate, whereas 1NMPP1-treated zygotes arrested prior to sporulation (Figure S5D). However, flow cytometry analysis revealed that the 1NM-PP1 treatment did not prevent DNA replication in these zygotes, which arrested with 4C DNA content (Figure S5E). These zygotes also did not display mating behaviors. As alternative strategy to block zygotes in G1, we used *cycΔ5* zygotes, of which the majority fails to enter S-phase (Figure 5I). Remarkably, mating of *cycΔ5* gametes, which are elongated likely due to an extended cell cycle, produced a noticeable fraction of zygotes that grew a shmoo and underwent re-fertilization (~4% of zygotes; Figures 6E-F and Movie 12). We conclude that rapid exit from the G1 phase is required to prevent zygotic mating and re-fertilization, even in cells with intact zygotic signaling. These data are consistent with the view that Mei3 blocks zygotic mating and re-fertilization by promoting the premeiotic S-phase.

### Mei2 blocks re-fertilization independently of cell cycle progression

While the role of Mei3 in blocking re-fertilization can be explained by its function in promoting the premeiotic S-phase, it remained unclear whether Mei2 blocks re-fertilization solely by promoting cell cycle progression or through other means. A role in promoting the meiotic cell cycle is suggested by our data and previous work (Watanabe and Yamamoto, 1994), as some *mei2Δ* cells arrest in G1 (Figure 6A). In addition, while *mei3Δ pat1Δ mei2Δ* triple mutant zygotes arrest in G1, re-expression of *mei2* in these cells (in the *SMS* background) restores a quasi-normal meiotic cycle (Figure 5). However, the higher frequency of lysis and re-fertilization observed in *mei3Δ mei2Δ* compared to *mei3Δ* zygotes, which are both arrested in G1, also indicates a cell cycle-independent function of Mei2 in blocking zygotic mating. To probe this point further, we compared G1-arrested *cycΔ5 mei3Δ mei2Δ pat1Δ* zygotes with or without Mei2 expression from the *p*^*mei3*^ promoter. Expression of Mei2, which in these G1-arrested zygotes is devoid of Pat1 inhibition, strongly repressed zygotic growth, mating, lysis and re-fertilization (Figure 7A-C and Movie 13). Consistently, the Scd2-GFP polarity patch was not present after fusion necks expanded in mutant zygotes expressing Mei2 (Figure 7D and Movie 14). We conclude that Mei2-dependent mating blocks largely operate independently of cell cycle progression.

**Figure 7.**
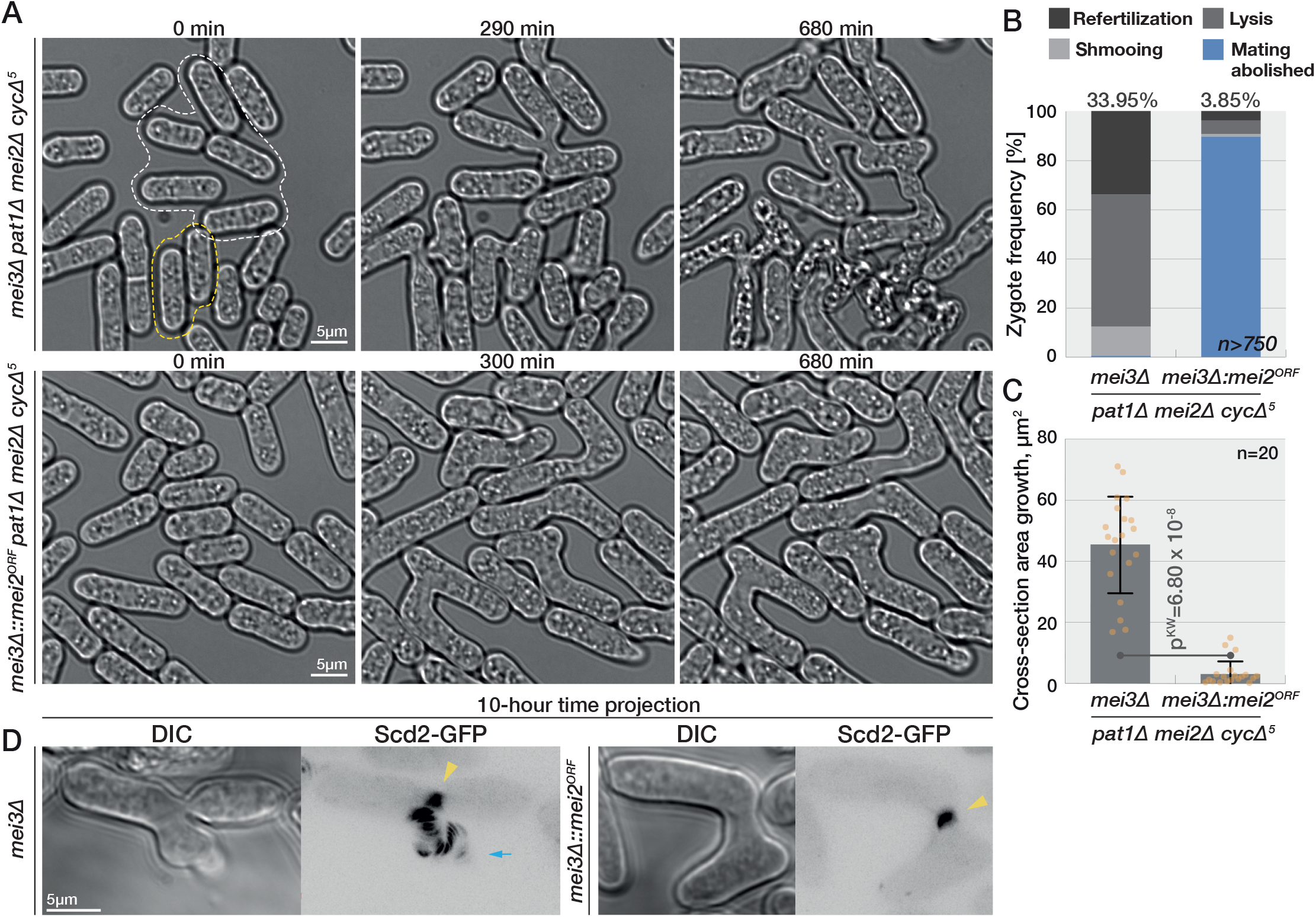
Mei2 blocks zygotic mating independently of cell cycle progression. **(A)** Mating mixtures of *SMS cycΔ5* cells (bottom), which induce Mei2 post-fertilization, and control cells that lack Mei2 (top). Mei2 expression prevents zygote lysis (yellow outlines) and re-fertilization (white outlines). **(B)** Quantification of phenotypes in zygotes as in (A). **(C)** Quantification of growth in zygotes as in (A) in the 10h after concave neck expansion was completed. Dots show individual measurements, bars mean values and error bars standard deviation. p-values for indicated comparisons were obtained from the Kruskal-Wallis test. **(D)** Projection of 60 timepoints during 10h from the time of fertilization of polarity and growth marker Scd2-GFP. The yellow arrowheads point to the signal at the neck. Scd2 is restricted to the neck in the zygote expressing Mei2 (right) but continuously trails the growth projection in the zygotes lacking Mei2 (blue arrow).

## Discussion

Zygote formation is a key developmental transition, where the newly formed diploid cell has to exit the mating program and re-enter the cell cycle. Our previous work established that the Mei3 and Mei2 proteins play a critical role in preventing zygote re-fertilization. These two proteins are part of a well-characterized signaling cascade that induces meiosis, where Mei3 relieves the inhibition of Mei2 by the kinase Pat1. Whereas most studies on meiosis have used data obtained from cells with inactivated Pat1 or from diploid strains that are unstable, here we tracked cells throughout the natural sexual lifecycle and performed detailed phenotypic analysis of mutants lacking Mei3, Pat1 and/or Mei2. Our long-term imaging combined with flow cytometry show that zygotes with defective signaling not only fail to initiate meiosis, but exhibit cell cycle defects and aberrant sex that results in re-fertilization and cell lysis. In contrast to the proposed linear meiotic signaling, we establish that Mei3 and Mei2 play independent functions to prevent re-fertilization and cell death. Mei3 serves to promote premeiotic DNA replication and thereby brings the zygote out of the mating-permissive G1 phase. Mei2 repression of mating behaviors is largely independent of both the cell cycle progression and activation by Mei3. Together these effects synergize to repress mating and confer the zygotic state (Figure 8).

**Figure 8.**
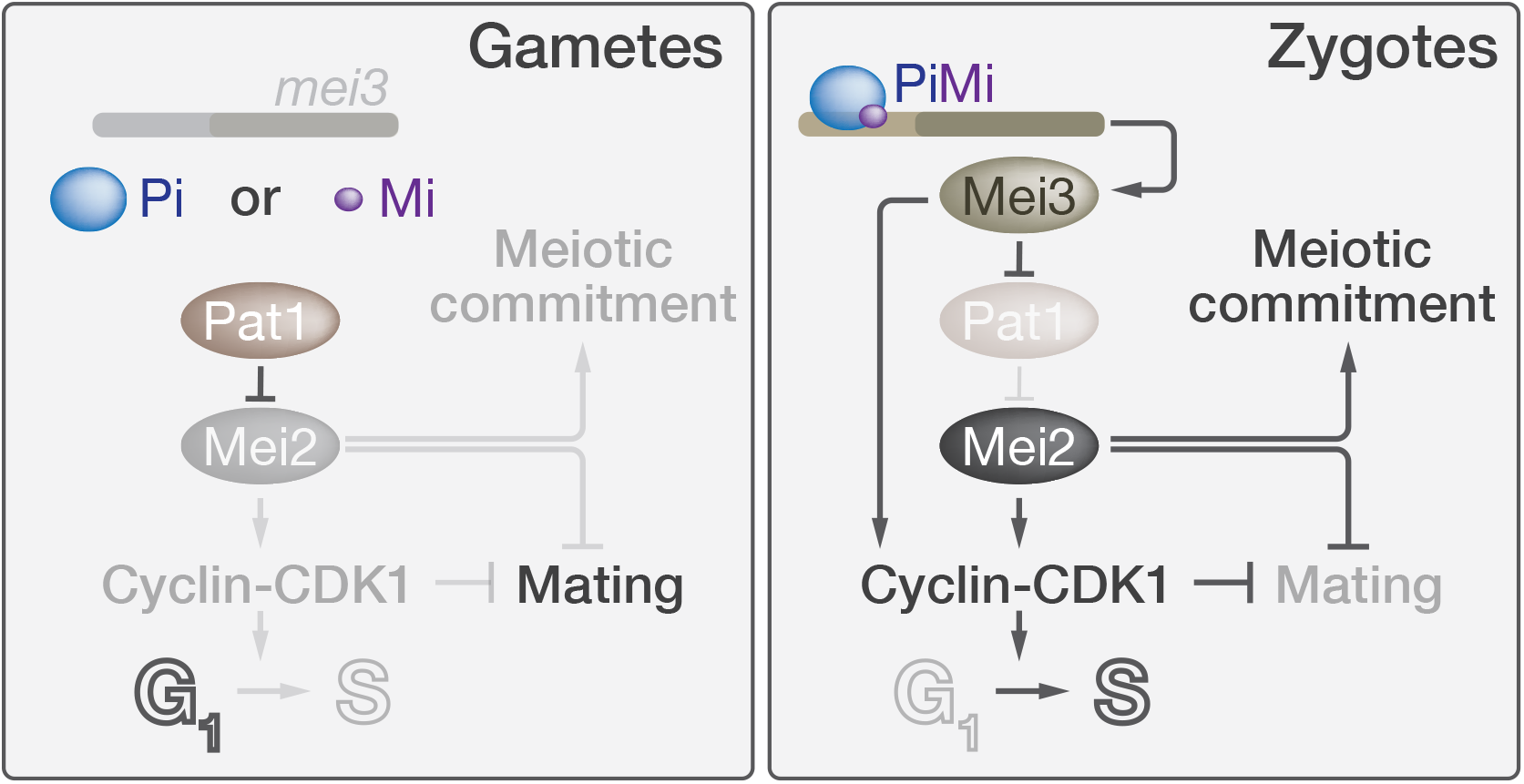
Model for gamete to zygote transition in fission yeast. In G1-arrested gametes (left panel), which express either Pi or Mi, *mei3* is not expressed and Pat1 maintains Mei2 repression to prevent meiosis and allow mating. Post-fertilization (right panel), the Pi-Mi complex induces Mei3 expression which relieves Mei2 inhibition. Mei3 and Mei2 independently activate CDK1 to repress mating and drive the G1-exit. Additionally, Mei2 commits zygotes to meiosis and imposes a second, cell cycle-independent block to mating.

### Re-fertilization blocks protect zygotes from death

Both Mei3 and Mei2 strongly contribute to preventing zygote re-fertilization, with 10-15% of *mei3Δ mei2Δ* double mutant zygotes fusing with an additional gamete, consistent with our previous observations (Vještica et al., 2018). However, an even more prevalent phenotype of *mei3Δ mei2Δ* zygotes is cell lysis. To test whether lysis may be due to failed zygotic fusion attempts, we developed a tool to target proteins for degradation specifically in zygotes. By linking a E3 ligase domain to the GFP- or mCherry-binding proteins, which bind their target with nanomolar affinities, this tool can be used to target any GFP- or mCherry-tagged protein for degradation. While combination of degron and target in the same cell leads to constitutive target degradation, as we have shown for cytosolic GFP and mCherry, the mating of gametes carrying the degron for the target expressed in the partner cell leads to target degradation specifically in zygotes. This is particularly interesting to test the zygotic role of essential proteins or proteins necessary for the earlier mating steps, and thus we expect it will be useful beyond the specific question asked here.

Using this zygotic degradation tool, we now show that the cell death of *mei3Δ mei2Δ* zygotes depends on the formin Fus1, the critical mating-specific organizer of the actin fusion focus. Indeed, degron-induced Fus1 depletion reduced fusion in gametes and lysis of *mei3Δ mei2Δ* zygote to similar extent, suggesting that most if not all lysis is due to erroneous activity of Fus1. This shows that zygotic death is due to ectopic formation of the fusion focus, which promotes local cell wall digestion. While we don’t formally show it, these ectopic fusion attempts are most likely due to zygotes becoming autocrine cells, which exhibit similar lysis phenotypes (Dudin et al., 2016), as fertilization brings together the production and perception machineries for both pheromones. We conclude that post-fertilization mating blocks not only prevent polyploid formation to ensure genome maintenance across generations, but also are essential to prevent zygotic death.

The high rates of re-fertilization and cell death in *mei3Δ mei2Δ* zygotes point to strong selective pressures to evolve mating blocks. In principle, species where sex relies on a diffusible signal produced by one and perceived by the other partner will lead to autocrine signaling post-fertilization. Thus, we expect that mechanisms to downregulate autocrine signaling and prevent cell death post-fertilization may be a common feature of sex in many fungi (Wallen and Perlin, 2018) and possibly other walled organisms that employ diffusible signals (Frenkel et al., 2014; Luporini et al., 2016).

### Mei3 promotes S-phase entry independently of Mei2

The strong phenotype of *mei3Δ mei2Δ* zygotes is a combination of the contributions of Mei3 and Mei2 functions, whose individual deletion produce much milder phenotypes. Multiple lines of evidence indicate that, in addition to its role in relieving Pat1 inhibition of Mei2 (Li and McLeod, 1996), Mei3 drives S-phase entry. First, *mei3Δ* and *mei3Δ mei2Δ* but not *mei2Δ* zygotes are blocked before S-phase, consistent with previous work on *mei3* (Egel and Egel-Mitani, 1974). Second, overexpression of Mei3 prevents the G1-arrest in response to nitrogen starvation. Third, Mei3 expression advances the premeiotic S-phase entry in *SMS* zygotes, which lack *pat1* and induce Mei2 expression only post-fertilization. Indeed, in the aforementioned experiments, the *pat1* gene is dispensable for the Mei3-driven cell cycle progression and thus Mei3 must operate independently of Mei2 activation. The weak effect of *mei3* deletion in *SMS* compared to otherwise WT zygotes indicates a role of Mei2 in pushing the cell cycle (see below), which Mei3 also contributes to by relieving Pat1 inhibition of Mei2. We conclude that an important role of Mei3, independent of its activation of Mei2, is to promote S-phase entry.

While the initial studies on meiosis suggested that Mei2 is essential for premeiotic S-phase (Watanabe and Yamamoto, 1994), these have relied on h+/h-diploids, which are unstable. We have revisited this work and introduced controls that ensure we are analyzing the h+/h-diploids stimulated by pheromone signaling. Our results mirror those obtained from zygotes and find that S-phase entry completely relies on Mei3 and only partially on Mei2. A direct implication of this result is that initiation of meiosis upon Pat1-inactivation, a widely used tool to initiate azygotic meiosis (Cipak et al., 2014), does not fully reproduce the physiological process. Instead, azygotic diploids with inactivated Pat1 are analogous to *SMS* zygotes as both lack Mei3 and Pat1 functions and start S-phase with a delay. This conclusion agrees with other notable differences between Pat1 inactivation-induced meiosis and normal zygotic meiosis (Bähler et al., 1991; Cipak et al., 2014) and suggests that Mei3 (and possibly Pat1) may have additional roles to that of regulating Mei2 activity.

One important question for the future is how Mei3 promotes premeiotic S-phase. A detailed recent study on the meiotic roles of cyclins found a number of differences between zygotic and azygotic meiosis (Gutiérrez-Escribano and Nurse, 2015). In particular, removal of all non-essential cyclins almost fully impaired S-phase entry during azygotic meiosis, but only partially prevented zygotic sporulation. Consistent with this work we found that the deletion of non-essential cyclins impairs the progression through premeiotic S-phase. As the G1-block is exacerbated in *cycΔ5* zygotes lacking Mei3 (but expressing active Mei2), we hypothesize that Mei3 at least in part promotes the activity of the only remaining cyclin Cdc13-CDK complex.

### Cell cycle progression acts as zygotic mating block

Irrespective of the precise molecular mechanisms by which Mei3 promotes premeiotic S-phase, this role is critical to prevent zygotic mating. Indeed, three lines of evidence show that cell cycle progression prevents mating post-fertilization. First, mating of *mei2Δ* zygotes correlates with their cell cycle stage, with only G1 and early S phase zygotes displaying mating behaviors. Second, G1 arrest upon removal of non-essential cyclins leads to re-fertilization, even in presence of functional Mei3-Pat1-Mei2 signaling. Third, forced cell cycle progression by zygotic expression of the G1 cyclin Puc1 substantially decreases the mating behaviors of *mei3Δ mei2Δ* zygotes. The observation that Puc1 reduces mating even in cells that fail to replicate their genomes suggests that CDK represses mating independently of the cell cycle transition. These data are in line with transcriptional genome-wide studies of meiosis showing that the expression of key mating regulators rapidly declines as cells progress through the meiotic cell cycle (Mata et al., 2002). They also agree with the well-established knowledge that gamete mating strictly requires G1-arrest, as CDK activity inhibits Ste11 at both pre- and post-transcriptional levels (Kjaerulff et al., 2007; Shimada et al., 2008). Antagonism between CDK activity and pheromone signaling is also well-established in other fungal organisms, such as *S. cerevisiae* (Kemp and Sprague, 2003; Peter and Herskowitz, 1994; Strickfaden et al., 2007). Thus, we propose that one important way Mei3 blocks re-fertilization is by promoting CDK activity.

### Mei2 blocks zygotic mating independently of cell cycle progression

The role of Mei2 is more complex, with at least three different functions. First, Mei2 is absolutely critical to induce meiosis. Indeed, we found that a fraction of *mei2Δ* zygotes enter a cell cycle akin to mitosis, with assembly of a single microtubule spindle followed by formation of a cytokinetic ring and septation. This indicates that upon cell fusion, zygotes lacking *mei2* can re-enter the cell cycle (driven by Mei3), but that this cycle is mitotic. Thus, Mei2 is the master specifier of meiosis. Second, Mei2 contributes in driving the meiotic cycle forward. Indeed, a fraction of *mei2Δ* zygotes and azygotic diploids are blocked or delayed in G1-phase. The ability of Mei2 to drive meiosis in SMS zygotes lacking Mei3 and Pat1 also shows that active Mei2 is in principle sufficient to allow a whole meiotic cycle, albeit with a G1 delay. A third function of Mei2, independent of promoting the meiotic cycle, is to block mating in zygotes. Indeed, *mei3Δ mei2Δ* double mutants show substantially more re-fertilization, growth, and lysis than *mei3Δ* zygotes, even though both are blocked in G1. This is also evident form the difference in dynamics of the polarity regulators Scd2 and Myo52, which were relatively short-lived in *mei3Δ* but stable in *mei2Δ* zygotes. These dynamics differences are reminiscent of dynamics observed in gametes where low levels of pheromone lead to unstable polarity zones which stabilize at elevated pheromone levels (Bendezú and Martin, 2012), suggesting that Mei2 suppresses pheromone signaling or downstream response. Expression of Mei2 in *SMS* zygotes lacking all non-essential cyclins was also sufficient to block most mating behaviors including Scd2 patch formation. We conclude that Mei2 imposes the zygotic fate, characterized by meiotic competence and mating blocks.

Our results challenge the current understanding of Mei2 regulation. The canonical view is that Mei2 is kept inactive by Pat1 until Mei3 relieves this inhibition. However, the observation that *mei3Δ* mutants do not lyse post-fertilization like *mei3Δ mei2Δ* zygotes indicates that Mei2 must prevent the pheromone from stabilizing mating and fusion machineries even prior to Mei3-dependent activation. This suggests that Pat1 inhibition of Mei2 is not complete, a view in agreement with a previous report that Mei2 promotes efficient mating when Mei3 is not expressed (Otsubo et al., 2014). These findings raise the question of how Mei2 allows mating prior to fertilization but prevents mating-induced lysis post-fertilization. The simplest explanation may be that there are additional differential regulations of Mei2 between gametes and zygotes, for instance inhibition by TOR kinase (Otsubo et al., 2014) or other unidentified factors.

### Synergistic action of two mating blocks in zygotes

We propose that the dramatic mating phenotypes of *mei3Δ mei2Δ* double mutant zygotes are due to synergistic actions of Mei3 and Mei2, which act in parallel to block mating. Mei2 is the key inducer of the zygotic fate, which represses mating and promotes meiosis. Mei3 involvement is twofold: On the one hand, Mei3 promotes Mei2 activation and in this way acts upstream of Mei2 to promote meiosis and inhibition of the pheromone response. On the other hand, Mei3 promotes the exit from the mating permissive G1 phase, independently of Mei2, and in this way acts synergistically with Mei2. Indeed, we found that combining mutations in non-essential cyclins, which block zygotes in G1, with *mei2Δ* also show synergistic mating block defects. Prolonging the G1-arrest exacerbates the mating block defects of *mei2Δ* zygotes by allowing extra time to grow mating projection and thus increasing chances of re-fertilization and lysis. Collectively our work shows that in fission yeast, distinct zygotic signaling branches initiate cell cycle-dependent and independent re-fertilization blocks that ensure survival and ploidy maintenance.

Robustness of biological systems often relies on parallel mechanisms regulating a single process (Stelling et al., 2004). This is also the case for re-fertilization blocks in several organisms (Gilbert, 2000). For example, in amphibians, changes in the plasma membrane potential provide the so called “fast” block to re-fertilization, while secretion of an outer cell layer presents a physical barrier to further sperm entry and a “slow” block to re-fertilization (Iwao, 2014). Subsequent development in all instances requires cell cycle progression, but its relevance as a re-fertilization block has not been studied. Fungal sexual reproduction might have experienced similar evolutionary pressures for rapid and long-term blocks to re-fertilization and lysis. Thus, Mei2 and Mei3-dependent mating blocks may act analogously, as rapid and delayed blocks, respectively. The fertilization induced exit from G1 occurs only ~35 min after cell fusion, leading to the speculation that evolution could have selected for additional rapid re-fertilization blocks. Furthermore, the G1-exit alone may not be sufficient to fully block mating since gametes arrested in S-phase do mate, albeit at a very low frequency (Kjaerulff et al., 2007). As the Mei2-dependent mating block is already operational independently of Mei3 at the time of fertilization, it may act as a “fast” block preventing re-fertilization during this permissive time frame.

## Materials and methods

### Strains and genetic markers

All **strains** used in the study are shown in Table S1.

All **genetic markers** with detailed description are available in Table S2. As indicated, previously reported markers were either present in the lab stock, received as a gift, obtained from the National BioResource Project (yeast.nig.ac.jp/yeast/top.jsf) or fission yeast deletion library produced by Bioneer (Daejeon, Republic of Korea).

For markers generated in this study we provide sequences of plasmids and primers used to make them in the **Supplemental Sequences** on Figshare (https://figshare.com/articles/dataset/Source_Data_for_Vjestica_et_al_2020/12674219). Briefly, we used SIV vectors (Vještica et al., 2020) to integrate artificial constructs into the *ade6, his5, lys3 or ura4* genomic loci. The pRIP vector (Maundrell, 1993) was used to integrate GFP under the *p*^*mam2*^ promoter at the *mam2* locus. Recombination assay markers were integrated into either *leu1*-*apc10* or *aha1-SPBC1711.09c* intergenic regions. Artificial degrons were integrated at either *ura4* or *fus1* loci. Fluorophore tagging of fission yeast proteins was performed at their native genomic loci with the exception of the S-phase marker Pcn1 which was expressed as second copy. To achieve constitutive gene expression, we used the strong promoters of *act1* and *tdh1* genes or moderate promoters of the *pcn1* gene or the SV40 viral promoter. To induce mating type-specific expression of genes we used either the M-cell specific *p*^*mam1*^ and *p*^*mam2*^ promoters or the P-cell specific *p*^*map3*^ promoter. Zygote-specific gene expression was achieved by using the *p*^*mei3*^ promoter. All constructs were sequenced and their integration into the genome confirmed by genotyping.

To prevent the *mat1* locus switching, wild-type H1-homology box was replaced with the one amplified from the PB9 strain carrying the ***H1Δ17* mutation**(Arcangioli and Klar, 1991; Vještica et al., 2018).

The **DegRed** artificial degron was comprised of the N-terminus of E3 ligase Pof1 (a.a. 1-261) fused to the mCherry-binding protein (ChBP, Fridy et al., 2014). Similarly, the **DegGreen** construct consisted of the Pof1 N-terminus (a.a. 1-261) fused to the GFP-binding protein (GBP, Rothbauer et al., 2006). Both constructs were placed under the regulation of the strong *p*^*tdh1*^ promoter (1000bp upstream the START codon) and the budding yeast ADH1 transcriptional terminator and integrated at either the *ura4* or *fus1* locus. We note that the two artificial degrons differ significantly in the ability to deplete target proteins (Figure 2B, 2C) for reasons that are not immediately clear. It is possible that the degron activities differ due to different expression levels, strength of nanobody-target binding or availability of residues for ubiquitination. We highlight that DegRed and DegGreen lack the WD40 repeats that target Pof1 to native target proteins and that neither caused any obvious defects in cells.

### Growth conditions

Note that all strains used in the study are prototrophs. The growth conditions used for experiments with **haploid cells** are detailed in (Vjestica et al., 2016). Briefly, freshly streaked cells were inoculated into MSL +N medium and incubated overnight at 25°C with 200 rpm shaking to exponential phase. The next day cultures were diluted to OD_600_ = 0.025 in 20 ml of MSL +N medium and incubated overnight at 30 °C with 200 rpm agitation to exponential phase. Experiments on exponentially growing cultures were performed at this point. For analysis of mating cells, homothallic cells or 1:1 mixtures of heterothallic cells were pelleted for one minute at 1000g, shifted to microcentrifuge tubes and washed three times with 1 ml of MSL-N medium. To prepare the mating mixtures for **flow cytometry** and **quantifications of sporulation, fusion and mating efficiencies**, we resuspended 1 OD_600_ (~1.4 × 10^7^ cells for our spectrophotometer) in 20μl of MSL -N media, dropped them onto solid MSL -N media and incubated at 25°C for 24h unless otherwise indicated. For **time-lapse imaging** and related quantifications, we diluted cells in 3 ml of MSL-N medium to final OD_600_ = 1-1.5 and incubated them at 30 °C with 200 rpm agitation for 4–6 h. The exception is the cycΔ5 mutants, which once nitrogen starved rapidly formed clumps difficult to image. Thus, we either shortened the incubation in liquid MSL -N to 2-3h, or alternatively, starved the *h+* and *h-* cells separately for 5 hours before mixing them immediately prior to imaging. Finally, cells were mounted onto MSL-N agarose pads, covered with a coverslip and the chamber was sealed using VALAP (vaseline, lanolin and paraffin; 1:1:1). We directly proceeded with imaging, or incubated slides until cells were 24h without nitrogen before scoring for phenotypes. The same growth conditions were used in **nitrogen starvation** experiments with a single heterothallic strain.

To summarize, we monitored exponentially growing cells in MSL +N liquid media and mating on MSL -N solid agar media plates or agarose pad chambers. We note that we observed quantitative differences between matings performed in different conditions (*e.g.* mating efficiency is higher on MSL -N solid agar plates) possibly due to limitation of nutrients and lower cell density in agarose chambers. Therefore, in all experiments, control strains were always run in parallel with identical conditions.

To obtain **diploid cells** through ***ade6* complementation**, we mated *ade6-M210* and *ade6-M216* strains on ME media and the next day streaked cells onto EMM media lacking adenine. As soon as visible, colonies of adenine prototrophs were restreaked and inoculated for experiments the following day. We paid attention that cells never overgrow as this may lead to meiotic activation. To obtain *H1Δ17* **diploids**, we mated on ME heterothallic strains that have the *H1Δ17* allele linked with different antibiotic resistance markers (Vještica et al., 2018). The next day we streaked cells onto YE media selecting for both antibiotics and waited for double resistant clones to form. We cultured both types of diploids starting with fresh cells OD_600_ = 0.025 in 20 ml of MSL +N media lacking amino acids. Cells were incubated overnight to OD_600_ = 0.3-0.5, collected by centrifugation at 1000g for 1 min, washed in MSL-N media three times. For **time-lapse imaging**, we imaged cells in the CellASIC ONIX microfluidic device (Merck Group, Darmstadt, Germany). For **flow cytometry and snapshot images**, we resuspended cells in 25 ml of MSL-N media to final O.D._600_ = 0.5. 5ml aliquots were collected at indicated timepoints, cells pelleted by centrifugation at 1000g for 1 min, supernatant discarded, and cell pellet thoroughly resuspended in 50 μl of MSL-N on ice before ethanol fixation.

To **evaluate the meiotic recombination frequency**, we prepared mating mixtures on solid MSL-N media as described above. After 48h incubation at 25°C, we collected cells using inoculation loops, resuspended them in 1ml of MSL-N media containing 10μl of glusulase (NEE154001EA, Perkin-Elmer) and incubated the samples for 24h at 30°C. Samples were then washed 3 times in water and we visually confirmed that glusulase treatment killed all cells except spores, thus ensuring that subsequent analyses evaluated only the filial generation. Samples were then resuspended in YE media and incubated at 30°C for ~8 hours. We diluted samples to final O.D._600_ = 0.025 and grew them to exponential phase for additional ~12 hours when samples were subjected to flow cytometry.

**Genetic crosses** were performed by mixing freshly streaked strains on either ME or MSL -N media. We note that MSL-N media enhanced mating and sporulation of mutants lacking multiple cyclins and zygotic signaling components.

Freshly streaked heterothallic strains carrying the **analogue sensitive***cdc2as-M17* alleles were pre-cultured separately in 3ml of MSL+N medium for 8-9 hours at 30°C with 200 rpm agitation before being diluted to OD_600_ =0.05 in 20 ml of MSL+N. The cell cultures were then incubated overnight at 18°C with 200 rpm rotation. The next morning, cells were washed 3 times in 20 ml of MSL-N before and resuspended in MSL-N OD_600_=1.5. Samples were split into two glass tubes with 3 ml in each and incubated at 18°C with 200 rpm agitation for 16 hours to arrest cells in G1-phase. For each strain culture, one tube was then treated with 25 μM **1NM-PP1**(Merck, Darmstadt, Germany, Cat No. 529581-1MG, 10 μM stock in DMSO) and the other tube with the control DMSO solvent. The cultures were incubated at 18°C for 1h and *h+* and *h-* cells mixed at a 1:1 ratio according to the treatment. Cells were further incubated at 18°C 200 rpm to allow mating.

We used standard conditions and protocols for genetic manipulations (Hagan et al., 2016). Briefly, yeast cells were grown in standard fission yeast YES media at either 25°C or 30°C and using 200 rpm rotators for liquid media. Cells were transformed with the lithium-acetate protocol (Hagan et al., 2016). For selection we supplemented YES with 100 μg/ml G418/kanamycin (CatNo.G4185, Formedium, Norfolk, UK), 100 μg/ml nourseothricin (HKI, Jena, Germany), 50 μg/ml hygromycinB (CatNo.10687010 Invitrogen), 100 μg/ml zeocin (CatNo.R25001, ThermoFischer), and 15 μg/ml blasticidin-S (CatNo.R21001, ThermoFischer). To our knowledge, this work is the first fission yeast report that employs ***patMX* dominant marker** to select for growth on glufosinate-ammonium. The *patMX* sequence was obtained from pAG31 plasmid (Goldstein and McCusker, 1999) and introduced into the pAV0892 (see Supplemental Sequences). After plasmid transformation into the prototroph yeast, cells were plated onto non-selective YES media and incubated overnight at 30°C. The next day we replicated the cells onto the EMM media without amino acids and supplemented with ~400 μg/ml of glufosinate-ammonium (CatNo. G002P01G, Cluzeau Info Labo, Sainte-Foy-La-Grande, France) and incubated them at 30°C. If background growth was observed after a couple of days, we replicated cells once more onto the fresh selective media. Individual colonies developed within 5 days.

### Flow cytometry

To prepare nitrogen starved and mated cells **for DNA content measurements**, we collected cells from solid media using inoculation loops and resuspended them in 1ml of MSL-N media pre-cooled to 4°C. Cells were pelleted by centrifugation at 1000g for 1 min at 4°C, supernatant discarded, and cells thoroughly resuspended in 50 μl of MSL-N on ice before ethanol fixation.

For **cell fixation**, we added 1ml of 70% EtOH (v/v in H_2_O) pre-chilled to −20°C and rapidly mixed by pipetting and vortexing for 5 sec, followed by incubation at −20°C for 15 min. Samples were once more vortexed for 5 sec, centrifuged at 4°C for 2 min at 2000g, supernatant discarded and cells gently resuspend in 1ml of room-temperature PBS (NaCl 8 g/l, KCl 0.2 g/l, Na_2_HPO_4_ 1.44 g/l, KH_2_PO_4_ 0.24 g/l, HCl adjusted pH = 7.4; filtered using 0.2μM filter). Samples were pelleted by centrifugation at room temperature for 1 min at 1000g, supernatant discarded, and cells resuspended in 1ml of PBS. We did not store the samples as we noticed that their quality degraded considerably already after a day at 4°C. Instead, we immediately **stained samples with Hoechst-33342**(Sigma-Aldrich B2261-25MG) at final concentration 25 ng/μl. We note that increasing the dye concentration produces a strong background signal. We resuspend 0.1 O.D._600_ of cells in 0.2 ml of PBS in 5ml flow cytometry Falcon tubes, added 1.8 ml of Hoechst working solution (27.7 ng/ml Hoechst-33342 in PBS) leading to final 25ng/μl Hoechst-33342 concentration and briefly vortexed. Tubes were closed and incubated in the in the dark ensuring thorough mixing using a rocking platform for 15-60 min before flow cytometry. We note that the signal produced by staining showed variation between experimental replicas, possibly due to stability of Hoechst-33342 stock (25mg/ml in DMSO, −20°C) and slightly different concentrations of the Hoechst working solution, which was freshly prepared for each experiment. Furthermore, experimental variation arose due to laser intensities and recording settings. Thus, we refrain from comparing samples from different experimental replicates and show in the same figure panel the data obtained from samples processed simultaneously.

**Flow cytometry was performed** on Fortessa instrument (Becton Dickinson, Franklin Lakes, USA) with proprietor software platform. The **analysis was performed** using the FlowJo (Becton Dickinson, Franklin Lakes, USA) software package. The 488 nm laser was used to determine the forward scatter (FSC) and side scatter (SSC). We used the 355 nm laser with 450/50 filter to detect Hoechst 33342 stained DNA, 488 nm laser with 505LP mirror and 550/50 filter to detect sfGFP and 561-nm laser line with 610LP mirror and 610/20 filter to detect mCherry.

**Raw data**(FCS format files) with sample legends and Source Data Figures detailing the **gating strategies** for each analysis are available from Figshare (https://figshare.com/articles/dataset/Source_Data_for_Vjestica_et_al_2020/12674219). Importantly, data presented in the same panel uses identical gates. We used FSC and SSC signal gating to exclude dust and clumped cells. Mating mixtures of P- and M-cells differentially labeled with sfGFP and mCherry provided fluorescent signals that were gated to distinguish gamete and zygote populations (Fig S3B), since gametes possess only one and zygotes both fluorophores. Importantly, these experiments also showed that gametes and zygotes produce populations with distinct FSC-SSC profiles (Figure S3B). We exploited this observation and used the FSC-SSC profiles to gate the zygote populations in mating mixtures produced by unlabeled gametes. When evaluating the frequency of meiotic recombinants, we used the red and green fluorescence signal to gate the populations of cells expressing neither, individual or both fluorophores.

### Microscopy and image processing

Images were obtained by wide-field microscopy performed on a DeltaVision platform (Applied Precision) composed of a customized inverted microscope (IX-71; Olympus), a UPlan Apochromat 100x/1.4 NA or 60x/1.4 NA oil objective, either camera (CoolSNAP HQ2; Photometrics) or camera (4.2Mpx PrimeBSI sCMOS camera; Photometrics), and a color combined unit illuminator (Insight SSI 7; Social Science Insights). Images were acquired using softWoRx v4.1.2 software (Applied Precision). We used the GFP-mCherry™ filterset to detect the green (Ex: 475/28, Em: 525/50) and red (Ex: 575/25; Em: 632/60) fluorescent proteins. We imaged a different number of Z-sections to best capture the structure of interest. We present either a single Z-plane a projection image.

All **image analysis and processing** was performed using Fiji software package (Schindelin et al., 2012) using either built-in modules or additional modules as specified. We used the MultiStackReg v1.45 module to adjust drifting time-lapses.

**Time-projection images** were prepared from 60 timepoints during 10h. Fluorescence channels show the maximum projection image. DIC images show either maximum or average projection image to allow for best visualization of the discussed phenotype.

### Quantifications and statistics

Sample sizes were not pre-determined. No randomization was used. No blinding was used. In analysis of imaging data, we excluded regions with multiple cell layers as we could not obtain reliable data. In morphometric analysis we excluded cells where we could not follow the shmoo growth because it went out of the imaging focal plane. We report p-values from Kruskal-Wallis test generated using Matlab function *ranksum()* and p-values from Welch’s t-test generated using Excel function *TTEST()*. Flow cytometry gating strategy is explained in detail and identical gates are applied to samples under comparison. All quantifications except those requiring long-term time-lapses were obtained from experimental triplicates. Variability between imaging chambers and their deterioration between different timepoints makes direct comparison between different imaging experiments difficult. For example, re-fertilization and lysis in meiotic signaling mutants increase over time and reach different incidence at 18hrs and 24hrs and thus results depend on when the chambers fail. Thus, we refrain from statistical comparison between different replicates and make comparisons only between samples processed and imaged simultaneously. We stress that we reached the same qualitative results and conclusions from experimental replicates. In panels 1B-C, 1F-G, S1C-D, 6E-F and S2A the wildtype was imaged separately from other samples.

Quantifications of **sporulation efficiencies** were performed on mating mixtures spotted on solid MSL-N media for 24h unless indicated otherwise. Samples from the edge of the mating mixture were scored for zygotic phenotypes by microscopy. We report percentages of each phenotype and p-values from Welch’s t-test.

Quantification of **zygotic phenotypes** was performed from DIC time-lapses where we could follow zygotes for 10 or more hours. We scored zygotes that underwent cell lysis, cell division, re-fertilization or neither and scored only the first observed event. Such scoring was adapted zygotes could carry out several events consecutively. For example, a zygote may undergo division and the daughter cells may then fuse with another partner. In quantifications that include of *mei3Δ* zygotes, we classified the remaining zygotes as non-dividing and refrain from denoting the percentage of shmooing since zygotes form subtle shmoos which we may underreport. Since *mei2Δ* and *mei3Δ mei2Δ* zygotes form prominent mating projections we additionally distinguish shmooing zygotes from those that abolished mating.

Quantification of **diploid phenotypes**(septation) was performed from DIC time-lapses.

In experiments targeting Fus1 with the synthetic degron, quantification of **fusion efficiencies** and **zygotic lysis frequencies** were performed from cells mating in agarose pad chambers 24h after nitrogen removal. Fusion efficiency was calculated as the percentage of fused gametes compared to the total gamete pairs formed. The detailed analysis of zygotic phenotypes was performed from long-term time-lapses.

DIC images were used for **analyses of zygotic growth and morphology**. We measured the **mating projection diameter** by determining the diameter of the half-circle that fitted in the shmoo tip. Our measurements correlated well with the diameter across the mating projections in *mei2Δ* and *mei3Δmei2Δ* zygotes, which could not be reliably measured for the subtle mating projections in *mei3Δ* zygotes. We compared the cell outlines immediately after the concave neck expansion ceases (initial perimeter) and ten hours later (final perimeter). Subtracting the area of two outlines determined the **growth of the cross-section area**. To quantify the displacement of the cell perimeter during this period we used the MorphoLibJ library (Legland et al., 2016). Briefly, for each zygote the initial perimeter was used to generate a binary image. We then used the MorphoLibJ function “Chamfer Distance Map” on the image to convert each pixel’s minimal distance from the initial outline into intensity of that pixel. We could then obtain intensities, which correspond to minimal distances from the initial perimeter, for each pixel on the final perimeter using built-in Fiji function “Plot Profile”. For each zygote we then determined the **maximum perimeter displacement** and the **coefficient of variation in perimeter displacement**(for perfect isometric growth, where all measured values are identical, coefV = 0). A minimum of 20 cells was analyzed and we report the mean measurement values and its standard deviation.

**Polarity patch lifetime** was determined from time lapses of cells co-expressing Scd2-GFP and Myo52-tdTomato imaged at a single plane with 10 min intervals. The patch was visually assessed and considered one entity when at consecutive timepoints the signal, irrespective of intensity, maintained position or moved to an adjacent location. Thus, we considered as a single patch those that trail the growth site, partially lose focus, transiently dim or slide across the cortex. We evaluated 10 or more cells per genotype and compared samples using the Kruskal-Wallis test.

Fluorescence quantifications to asses efficiency of the **artificial degron** was performed on average projections of images containing 12 z-sections with 0.5μm spacing. We measured the fluorescence in unlabeled wildtype cells, and the cells expressing either the fluorophore alone or together with the degron. We used the DIC channel to outline the cells, measured the mean fluorescence of 30 cells per sample and the slide background florescence. We subtracted the slide background fluorescence and compared samples using the Kruskal-Wallis test. We then calculated the average fluorescence for cells from each sample and its standard deviation. We set the average wildtype cell fluorescence to 0, the average fluorescence of cells expressing the fluorophore alone to 100 and used error propagation formula to calculate standard deviations.

Quantifications of **Mi induction** were performed on single Z-plane time lapses of nitrogen staved *H1Δ17* diploids. We used the DIC channel to outline the cells over time and measure mean cell fluorescence. We used these measurements to visually determine the time of Mi induction and align time-courses from different cells. The Mi signal also allowed us to visualize and outline the circular nuclei and measure the mean fluorescence. We then subtracted the slide background and report the mean fluorescence (line) and standard deviation from measurements of 8 cells.

Quantifications of the **S-phase timing** used the exchange of cytosolic mCherry between partner gametes to determine the fusion time (T_fusion_). Time of S-phase was recorded as the first timepoint with punctate GFP-Pcn1 localization. We analyzed only cells that fused within the first 5 hours since the onset of imaging since we noticed that zygotes formed at late timepoints took longer time to exit G1 in all crosses investigated, possibly due to depletion of nutrients from the agarose pads used for imaging. Additionally, we ensured that the populations we compare have similar average T_fusion_ and do not show statistically significant T_fusion_ differences (p^Kruskal-Wallis^ > 0.1).

To determine the **meiotic recombination frequencies**, in one parental strain we introduced at the *leu1* genomic locus sequences driving sfGFP expression from the strong *p*^*tdh1*^ promoter, and in the other parental strain at the *aha1* genomic locus sequences driving mCherry expression also from the strong *p*^*tdh1*^ promoter. We chose to integrate the constructs proximal to *leu1* and *aha1* loci in regions with no known function. Mating between these two parental strains yielded non-recombinant progeny that expresses one fluorophore and recombinant progeny that expresses either both or neither fluorophore. We used flow cytometry to determine the frequencies of recombinant and non-recombinant progeny and the genetic distance of the two loci in cM (centimorgan) was determined according to the following formula:

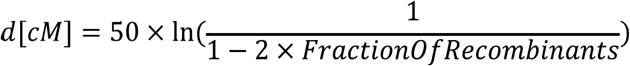

The experiment was performed in triplicate and we report mean values, standard deviation and p-values obtained using the Welch’s t-test. We obtained that the genetic distance between the *leu1* and *aha1* loci is 17.13±0.14 cM in wildtype crosses, which is consistent with previous report of ~18cM separating the *leu1* and *aha1*-proximal *his2* gene (Kohli et al., 1977; Saito et al., 2004).

## Data availability

The datasets generated and/or analysed during the current study are available from corresponding authors on reasonable request.

A single ZIP file (https://figshare.com/articles/dataset/Source_Data_for_Vjestica_et_al_2020/12674219) available from Figshare contains the Supplemental Sequences, quantification summaries and flow cytometry data distributed in subfolders corresponding to figure panels. Data used in quantification is presented in Microsoft Excel format. Flow cytometry data includes raw data files in the FCS format and Source data figures which show gating strategies in detail.

## Author contributions

AV and SGM conceived the project. MB performed and analyzed experiments shown in Fig S5D-E. GL provided input and constructs crucial for Fig 6C-D. LM assisted AV with data analysis in Fig 2 and Fig 7. PJN provided technical assistance. AV performed all other experiments and analyses. AV and SGM wrote the manuscript. SGM acquired funding.

## Acknowledgements

We thank Ingrid Billault-Chaumartin for advice and critical revising of the manuscript. We thank Viesturs Simanis for shared strains. This work was funded by an ERC consolidator grant (CellFusion) and a Swiss National Science Foundation grant (310030B_176396) to SGM.

## Supplemental Materials

**Table S1. Strains used in this study.**

Table S2. Description of all genetic markers introduced in strains in this study.

**Movie 1. Meiotic signaling mutants show distinct phenotypes.** DIC time-lapses show mating of meiotic signaling mutants. Note zygotes forming mating projections (yellow arrowheads) and *mei2Δ* zygotes dividing (red arrowhead)

**Movie 2. Division ring forms in***mei2Δ* **but not in***mei3Δ* **nor***mei3Δ mei2Δ* **zygotes.** Time-lapses show mating of meiotic signaling mutants. Fertilization is visualized as partner exchange of cytosolic mCherry (magenta) expressed from the P-cell specific *p*^*map3*^ promoter. Note *mei2Δ* zygotes (yellow outlines) that assemble an actomyosin ring labelled by Rlc1-sfGFP (green; arrowheads) and divide.

**Movie 3. Pat1 kinase is dispensable for division in***mei2Δ* **zygotes.** DIC time-lapses show mating of meiotic signaling mutants. Note that cell division (yellow arrowheads) occurs in *pat1Δ mei2Δ* but not in *mei3Δ pat1Δ mei2Δ* zygotes.

**Movie 4. Meiotic signaling mutants show distinct dynamics of growth and polarity markers.** Time-lapses show mating of cells expressing polarity and growth markers Scd2-GFP (green) and Myo52-tdTomato (magenta). The intense Myo52 signal formed at the time of fertilization (highlighted with white arrowheads in outlined cell pair) rapidly disappears in wildtype zygotes but persists (yellow arrowheads) in zygotes with impaired meiotic signaling.

**Movie 5. Zygote lysis is induced by aberrant fusion attempts.** Mating of *mei3Δ mei2Δ* gametes carrying the indicated halves of the Fus1 artificial degron system. Arrowheads point to zygotes with incomplete Fus1 artificial degron (top and middle panel) that maintain a strong Fus1-GFP (green) or Fus1-mCherry (magenta) signal post-fertilization. Zygotes with incomplete Fus1 degron undergo cell lysis (example marked with yellow outlines) more frequently than zygotes with the complete Fus1 degron (bottom panel).

**Movie 6. Mei3 promotes G1 exit in stable diploid cells.** Time-lapses show *H1Δ17* diploid cells expressing mCherry (magenta) and sfGFP (green) from P- and M-cell specific promoters *p*^*map3*^ and *p*^*mam1*^ after a shift to nitrogen-free media. Note septum formation (arrowheads) in *mei2Δ* mutant cells expressing both fluorescent proteins.

**Movie 7. The Pi-Mi complex formation precedes mating in diploid cells.** Time-lapses of *mei3Δ H1Δ17* diploid cells shifted to nitrogen-free media that express Mi-sfGFP and are either wildtype (top) or mutant (bottom) for Pi. Nuclear enrichment of Mi (red arrowheads) is Pi dependent and precedes the formation of mating projections (yellow arrowheads).

**Movie 8. Mei3 advances premeiotic-S phase entry.** Time-lapse shows DIC (grey) and the nuclear GFP-Pcn1 (green) in wildtype (top panels) and *SMS* zygotes (bottom panels). Fertilization is evident as partner exchange cytosolic mCherry (magenta) expressed in only one gamete. Punctate GFP-Pcn1 signal (arrowheads), indicative of S-phase, appears prior to karyogamy in WT but later in *SMS* zygotes.

**Movie 9. Zygotes lacking***mei3* **cannot progress to S-phase.** Time-lapses show DIC (left panels) and GFP-Pcn1 (right panels) during mating of wildtype and mutant cells. GFP-Pcn1 puncta (red arrowhead) form in wildtype, *mei2Δ*, *sme2Δ* and *mei4Δ* zygotes but not in *mei3Δ* and *mei3Δ mei2Δ* zygotes.

*Movie 10. Mating behavior correlates with the cell cycle* **phase in***mei2Δ* **zygotes.** Time-lapse of mating *mei2Δ* cells. Zygotes that persist in G1 (yellow outlines) form prominent mating projections (yellow arrowhead). Zygotes that progress through S-phase (blue outlines), evident from GFP-Pcn1 puncta (blue arrowhead), largely repress shmoo formation and instead divide.

**Movie 11. Forced cell cycle progression decreases mating behaviors of***mei3Δ mei2Δ* **zygotes.** Time-lapses show mating of control *mei3Δ mei2Δ* cells (top) and *mei3Δ mei2Δ* cells that express Puc1 from the zygote specific *p*^*mei3*^ promoter. GFP-Pcn1 (right panels) forms puncta (red arrowheads) in Puc1-expressing zygotes (blue outlines). The control G1-arrested zygotes form prominent mating projections and frequently lyse unlike Puc1-expressing zygotes which show little or no growth (yellow outlines) and reduced lysis.

**Movie 12. Zygotes lacking non-essential cyclins undergo re-fertilization.** DIC time-lapse shows mating of *cycΔ5* cells, which undergo re-fertilization (yellow outlines).

**Movie 13. Mei2 blocks zygotic mating independently of cell cycle progression.** DIC time-lapses show mating mixtures of *SMS cycΔ5* cells (right panel), which induce Mei2 post-fertilization, and control cells that lack Mei2 (left panels). Mei2 expression prevents zygote lysis and re-fertilization.

**Movie 14. Mei2 prevents zygotic growth.** Time-lapses show DIC (left) and Scd2-GFP (right) in mating mixtures of *SMS cycΔ5* cells (bottom), which induce Mei2 post-fertilization, and control cells that lack Mei2 (top). Note that the Scd2 signal dissipates after fusion in zygotes expressing Mei2 but trails the growth projection in the zygotes lacking Mei2.

---

**Figure S1.**
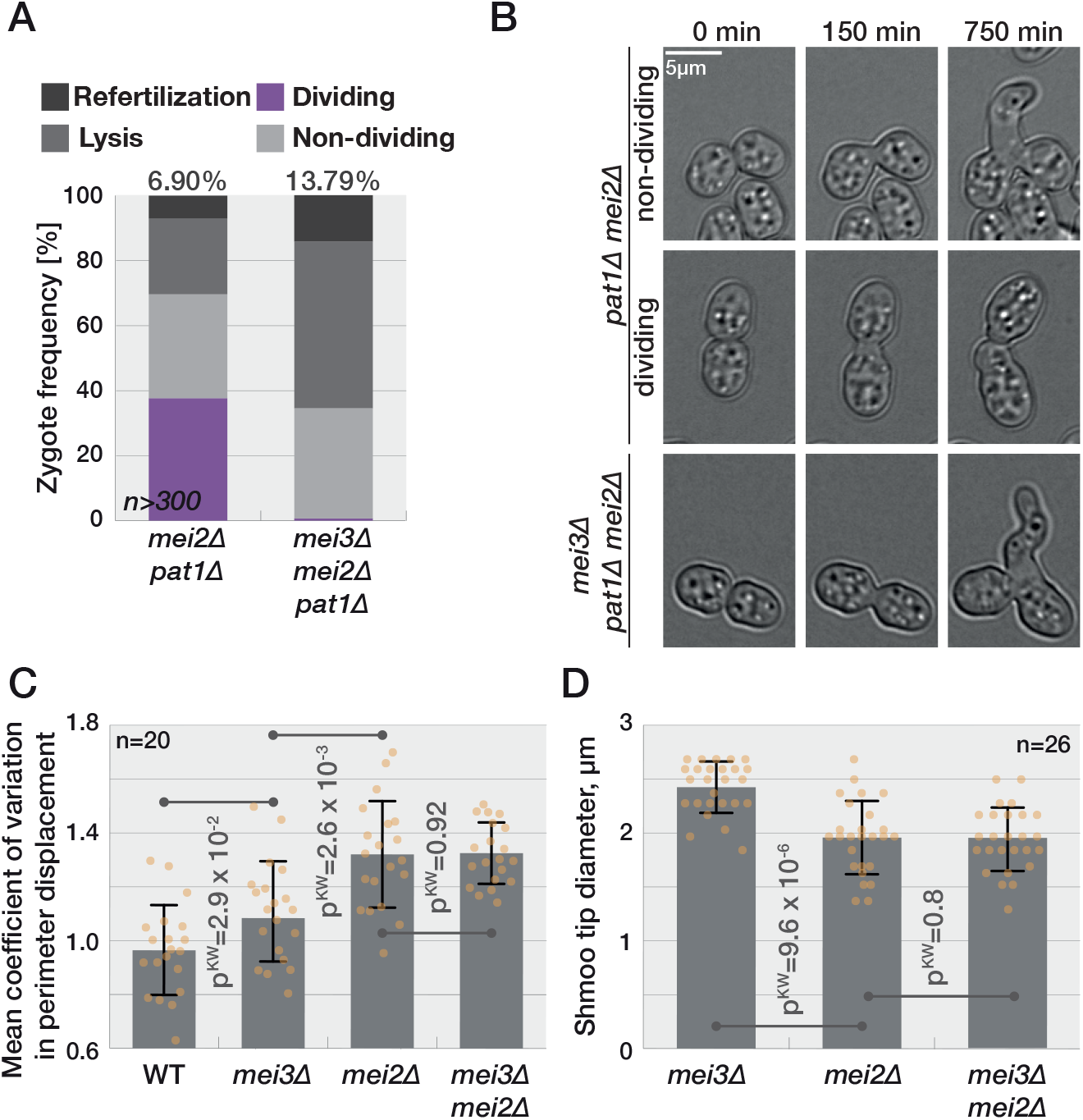
Mei3 has functions independent of Pat1 kinase. **(A)** Quantification of phenotypes observed by long-term DIC imaging of zygotes. Numbers at the top report frequency of re-fertilization. **(B)** DIC images of cells prior to partner fusion (0 min), at the time zygotes complete concave neck expansion (150 min) and 10h later (750 min). Note that a fraction of *pat1Δ mei2Δ* zygotes divides. **(C-D)** Quantification of zygotic growth observed by time-lapse imaging during 10h after neck expansion measured from outlines as shown in Fig 1C. Dots show individual measurements, bars mean values and error bars standard deviation. p-values for indicated comparisons were obtained from the Kruskal-Wallis test. In (C), we determined the zygote outlines at the time of completion of concave neck expansion and 10h later, which we then used it to determine the perimeter displacement for each pixel and quantify the coefficient of variation. Theoretically, perfect isometric growth shows zero coefficient of variation. The values obtained for the wildtype zygotes, which do not grow, are indicative of the measurement error. In (D) we quantified the diameter of the shmoos formed by zygotes.

**Figure S2.**
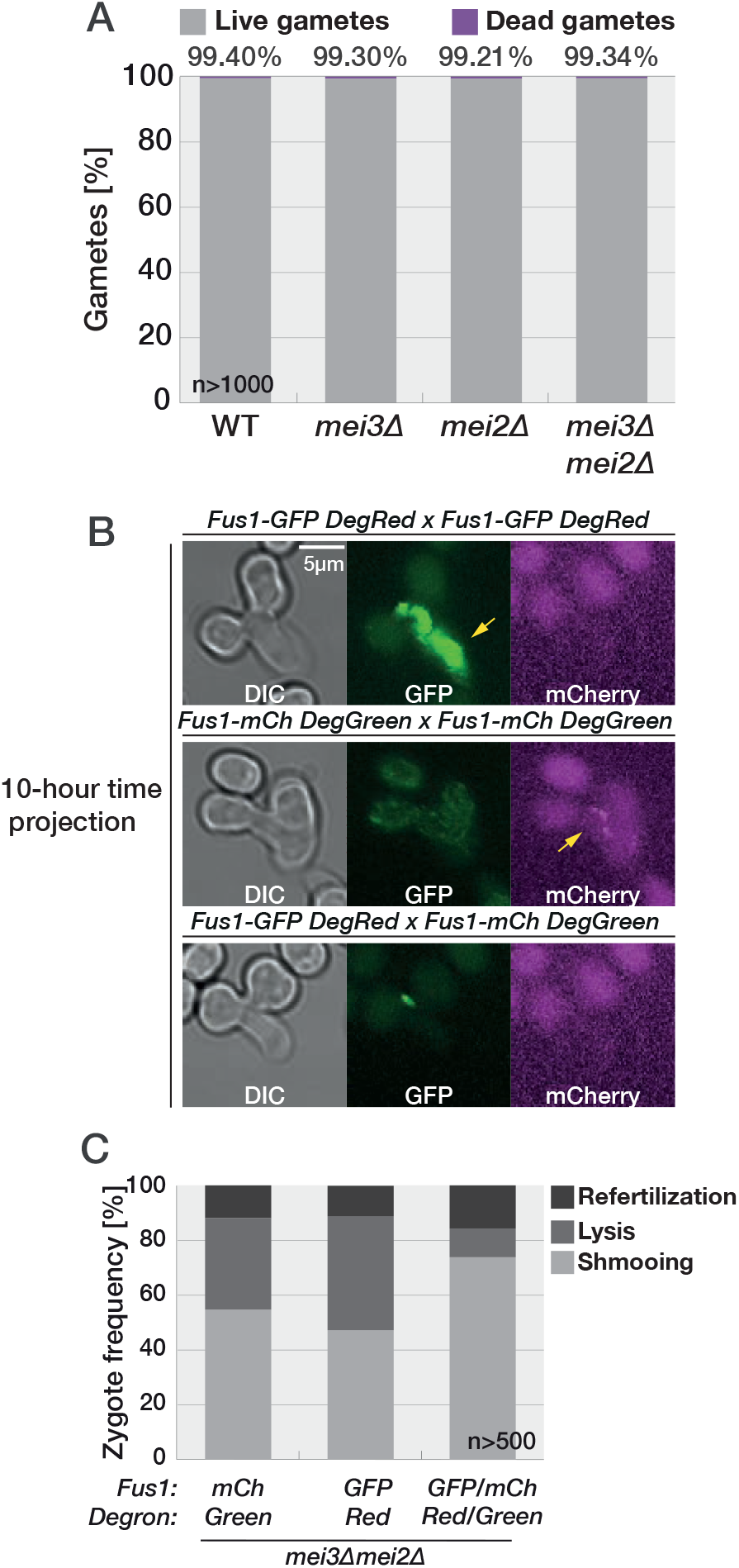
Zygote lysis is induced by aberrant fusion attempts. **(A)** The graph quantifies the frequency of lysed and intact gametes within 24h of mating as observed from DIC time-lapse imaging. Numbers on the top report the frequency of surviving gametes. *mei2Δ* and *mei3Δ mei2Δ* gametes do not lyse. **(B)** Projection of 60 timepoints obtained during 10h post-fertilization of *mei3Δ mei2Δ* gametes carrying the indicated halves of the Fus1 artificial degron. Zygotes with the incomplete Fus1 degron (top and middle panels) showed a high level of Fus1 fused to the fluorophore which trailed the growth projections (yellow arrows). Conversely, in zygotes that inherited a complete artificial degron from the two parents (bottom panel), Fus1 fluorescent signal was detected only at the time of fertilization followed by its rapid disappearance during growth of mating projections. Note that Fus1-mCherry is detectable post-fusion in *mei3Δ mei2Δ* zygotes that have an incomplete degron system but is not detectable in gametes due to low fluorescence signal. **(C)** Quantification of zygotic phenotypes observed during time-lapse imaging of mating between indicated *h+* and *h-* strains which produce zygotes with either incomplete (left and middle bar) or complete (right bar) artificial Fus1 degron. Note the decreased lysis in zygotes that carry the complete Fus1 degron system.

**Figure S3.**
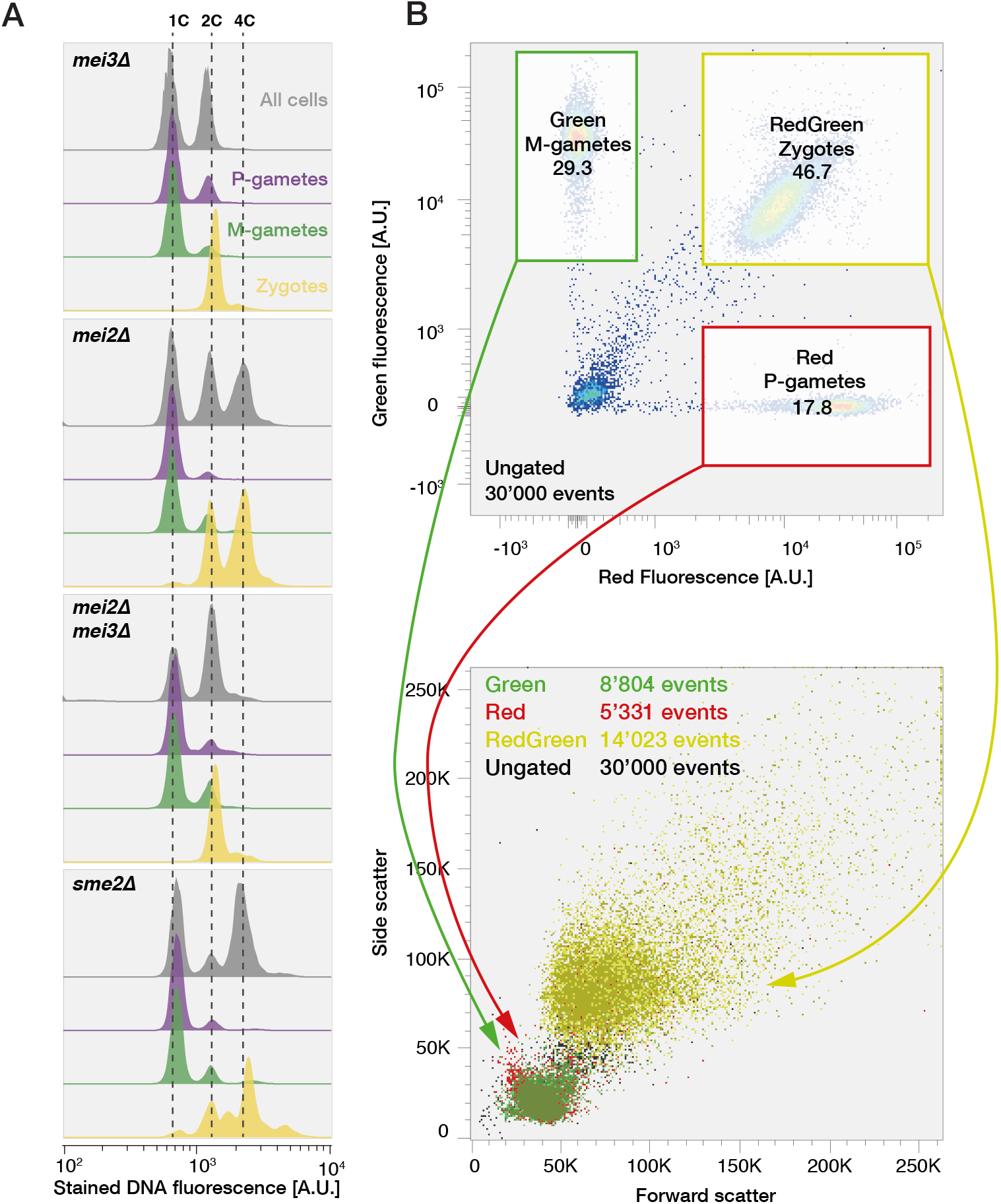
Mei3 promotes G1 exit and flow cytometry gating strategy. **(A)** Strategy used for flow cytometry during mating of homothallic strains which express sfGFP and mCherry from mating type specific promoters *p*^*mam*1^ and *p*^*map3*^, respectively. Haploid cells, which differentiate into either P- or M-gametes, induce a single fluorophore and thus can be distinguished from zygotes, which express both fluorophores. Flow cytometry analysis of Hoechst-stained DNA fluorescence (x-axis) 24h after transfer to medium lacking nitrogen (bottom). The y-axis shows the cell number normalized to mode. Profiles of all cells in the mating mixture (grey), gametes (green and magenta) and zygotes (yellow). Gametes largely arrest with unreplicated 1C genomes. *mei3Δ* and *mei3Δ mei2Δ* zygotes show unreplicated 2C content. *mei2Δ* and *sme2Δ* zygotes show a prominent 4C peak. **(B)** Flow cytometry analysis of mating mixtures produced between the *h-* strain expressing sfGFP and *h+* strain expressing mCherry from a constitutive promoter. Note that populations of gametes and zygotes gated according to red and green fluorescence (top panel) produce distinct populations in the forward and side scatter plot (bottom panel).

**Figure S4.**
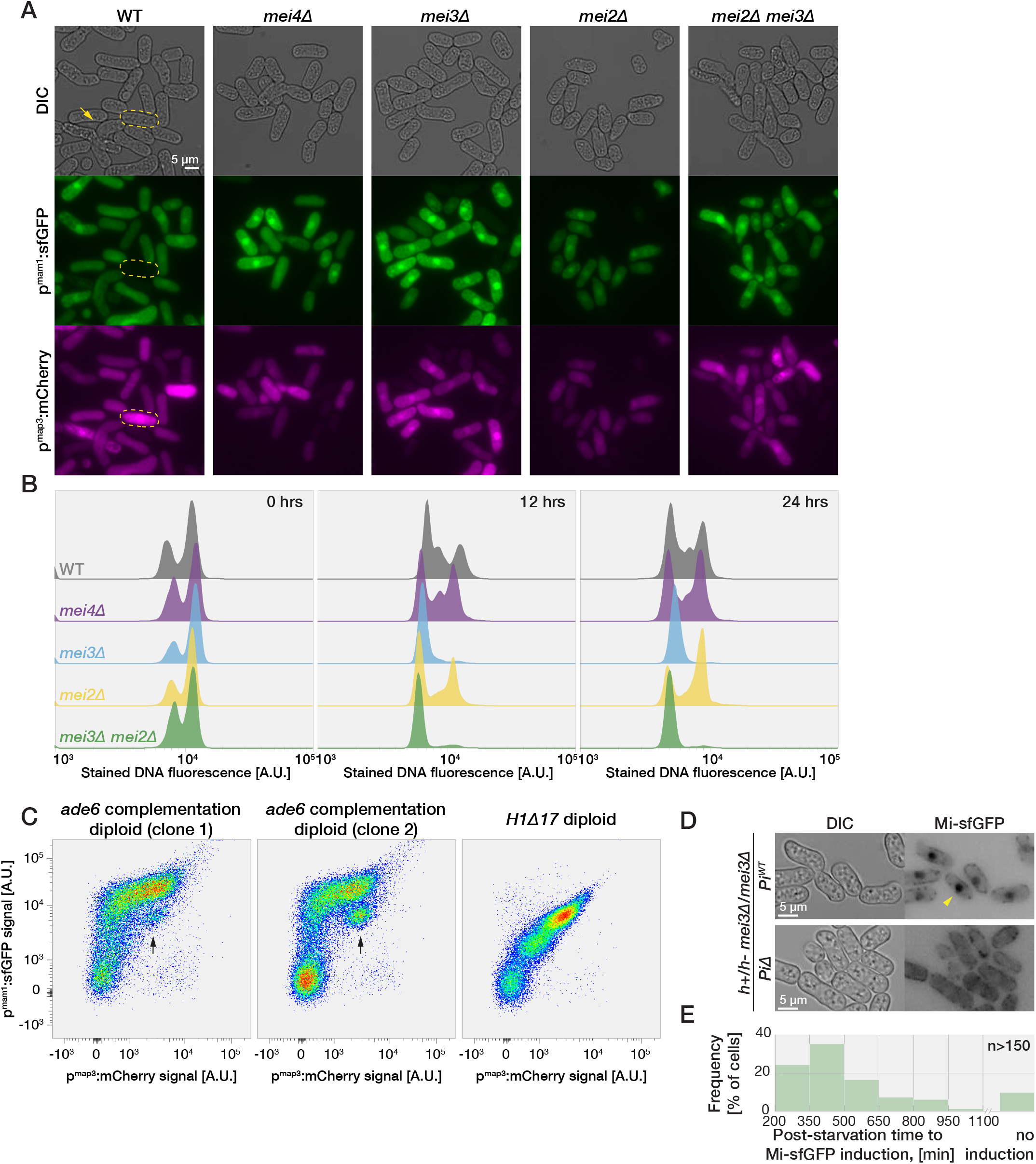
Mei3 promotes G1 exit in diploid cells obtained by *ade6* complementation. **(A)** Diploid cells obtained by *ade6* complementation expressing mCherry and sfGFP from P- and M-cell specific promoters *p*^*map3*^ and *p*^*mam1*^ 24h after removal of nitrogen. Note the mated pair (arrow) and a cell expressing only mCherry (yellow outline). **(B)** Flow cytometry analysis of Hoechst-stained DNA fluorescence (x-axis) in diploid cell used in (A) at indicated timepoints following nitrogen removal. The y-axis shows the cell number normalized to mode. Note that *mei3Δ* and *mei3Δ mei2Δ* mutants arrest with unreplicated genomes (2C) while wildtype, *mei4Δ* and *mei2Δ* cells replicate their DNA (4C). **(C)** Flow cytometry analysis of mCherry and sfGFP induction from the *p*^*map3Δ*^ and *p*^*mam1*^ promoters in two *ade6*-complementation diploids (left and middle panels) and a *H1Δ17* diploid (right panel) 24h after nitrogen removal. Note subpopulations of cells with high *p*^*map3*^ signal (arrow). **(D)** DIC and Mi-sfGFP images of nitrogen starved *mei3Δ* diploids which either express (top) or lack (bottom) Pi. Nuclear localization of Mi (yellow arrowhead) is dependent on Pi. **(E)** Quantification of the time required to detect Mi-sfGFP in diploid cells shifted to nitrogen-free media.

**Figure S5.**
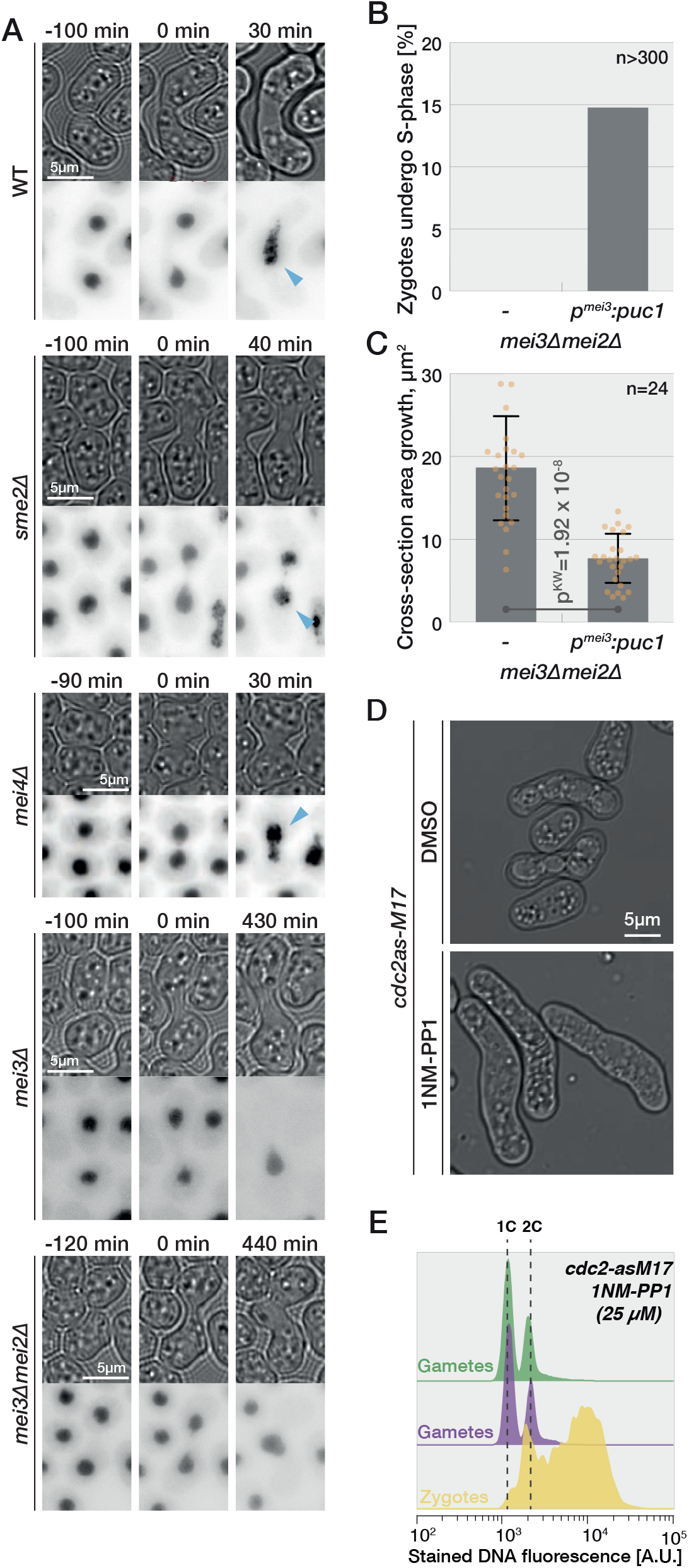
Cell cycle progression correlates with mating repression. **(A)** Time-lapses show DIC (top panels) and GFP-Pcn1 (bottom) during mating of wildtype and mutant cells. GFP-Pcn1 puncta (blue arrowhead) form in wildtype, *sme2Δ* and *mei4Δ* zygotes but not in *mei3Δ* and *mei3Δ mei2Δ* zygotes. **(B)** Quantification of zygotes passing through S-phase as observed from GFP-Pcn1 dynamics shown in Figure 6C. **(C)** Quantification of growth in G1-arrested zygotes in the 10h after concave neck expansion was completed. Dots show individual measurements, bars mean values and error bars standard deviation. p-values for indicated comparisons were obtained from the Kruskal-Wallis test. **(D)** DIC images of mated *cdc2as-M17* cells treated with 1NM-PP1 (right) or DMSO control (left). Mating projections are not observed in zygotes. **(E)** Flow cytometry analysis of Hoechst-stained DNA fluorescence (x-axis) in mating mixtures of *cdc2as-M17* cells. The y-axis shows the cell number normalized to mode. Gametes and zygotes were distinguished as in Figure 3B. Gametes largely arrest with unreplicated 1C genomes while zygotes show a prominent 4C peak.

## References

Aoi Y, Kawashima SA, Simanis V, Yamamoto M, Sato M. 2014. Optimization of the analogue-sensitive Cdc2/Cdk1 mutant by in vivo selection eliminates physiological limitations to its use in cell cycle analysis. Open Biol 4. doi:10.1098/rsob.140063

Arcangioli B, Klar AJ. 1991. A novel switch-activating site (SAS1) and its cognate binding factor (SAP1) required for efficient mat1 switching in Schizosaccharomyces pombe. EMBO J 10:3025–32.

Bähler J, Schuchert P, Grimm C, Kohli J. 1991. Synchronized meiosis and recombination in fission yeast: observations with pat1-114 diploid cells. Curr Genet 19:445–51. doi:10.1007/BF00312735

Beach DH, Klar AJ. 1984. Rearrangements of the transposable mating-type cassettes of fission yeast. EMBO J 3:603–10.

Beale KM, Leydon AR, Johnson MA. 2012. Gamete Fusion Is Required to Block Multiple Pollen Tubes from Entering an Arabidopsis Ovule, Current Biology. doi:10.1016/j.cub.2012.04.041

Bendezú FO, Martin SG. 2013. Cdc42 explores the cell periphery for mate selection in fission yeast. Curr Biol 23:42–7. doi:10.1016/j.cub.2012.10.042

Bendezú FO, Martin SG. 2012. Cdc42 Explores the Cell Periphery for Mate Selection in Fission Yeast. Curr Biol 1–6. doi:10.1016/j.cub.2012.10.042

Bendezú FO, Martin SG. 2011. Actin cables and the exocyst form two independent morphogenesis pathways in the fission yeast. Mol Biol Cell 22:44–53. doi:10.1091/mbc.e10-08-0720

Chang EC, Barr M, Wang Y, Jung V, Xu H-P, Wigler MH. 1994. Cooperative interaction of S. pombe proteins required for mating and morphogenesis. Cell 79:131–141. doi:10.1016/0092-8674(94)90406-5

Cipak L, Polakova S, Hyppa RW, Smith GR, Gregan J. 2014. Synchronized fission yeast meiosis using an ATP analog-sensitive Pat1 protein kinase. Nat Protoc 9:223–31. doi:10.1038/nprot.2014.013

Davey J, Nielsen O. 1994. Mutations in cyr1 and pat1 reveal pheromone-induced G1 arrest in the fission yeast Schizosaccharomyces pombe. Curr Genet 26:105–12. doi:10.1007/BF00313796

Dudin O, Bendezú FO, Groux R, Laroche T, Seitz A, Martin SG. 2015. A formin-nucleated actin aster concentrates cell wall hydrolases for cell fusion in fission yeast. J Cell Biol 208:897–911. doi:10.1083/jcb.201411124

Dudin O, Merlini L, Martin SG. 2016. Spatial focalization of pheromone/MAPK signaling triggers commitment to cell–cell fusion. Genes Dev 30:2226–2239. doi:10.1101/gad.286922.116

Egel R, Egel-Mitani M. 1974. Premeiotic DNA synthesis in fission yeast. Exp Cell Res 88:127–34. doi:10.1016/0014-4827(74)90626-0

Ekwall K, Thon G. 2017. Selecting Schizosaccharomyces pombe Diploids. Cold Spring Harb Protoc 2017:pdb. prot091702. doi:10.1101/pdb.prot091702

Frenkel J, Vyverman W, Pohnert G. 2014. Pheromone signaling during sexual reproduction in algae. Plant J 79:632–644. doi:10.1111/tpj.12496

Fridy PC, Li Y, Keegan S, Thompson MK, Nudelman I, Scheid JF, Oeffinger M, Nussenzweig MC, Fenyö D, Chait BT, Rout MP. 2014. A robust pipeline for rapid production of versatile nanobody repertoires. Nat Methods 11:1253–1260. doi:10.1038/nmeth.3170

Gilbert SF. 2000. Developmental biology, 6th Editio. ed. Sinauer Associates. https://www.ncbi.nlm.nih.gov/books/NBK10033/.

Goldstein AL, McCusker JH. 1999. Three new dominant drug resistance cassettes for gene disruption in Saccharomyces cerevisiae. Yeast 15:1541–53. doi:10.1002/(SICI)1097-0061(199910)15:14<1541::AID-YEA476>3.0.CO;2-K

Gutiérrez-Escribano P, Nurse P. 2015. A single cyclin-CDK complex is sufficient for both mitotic and meiotic progression in fission yeast. Nat Commun 6:6871. doi:10.1038/ncomms7871

Hagan I, Carr AM, Grallert A, Nurse P. 2016. Fission Yeast: A Laboratory Manual. Cold Spring Harbour Press.

Harigaya Y, Yamamoto M. 2007. Molecular mechanisms underlying the mitosis–meiosis decision. Chromosom Res 15:523–537. doi:10.1007/s10577-007-1151-0

Higuchi T, Watanabe Y, Yamamoto M. 2002. Protein kinase A regulates sexual development and gluconeogenesis through phosphorylation of the Zn finger transcriptional activator Rst2p in fission yeast. Mol Cell Biol 22:1–11. doi:10.1128/mcb.22.1.1-11.2002

Horie S, Watanabe Y, Tanaka K, Nishiwaki S, Fujioka H, Abe H, Yamamoto M, Shimoda C. 1998. The Schizosaccharomyces pombe mei4+ gene encodes a meiosis-specific transcription factor containing a forkhead DNA-binding domain. Mol Cell Biol 18:2118–29. doi:10.1128/mcb.18.4.2118

Imai Y, Yamamoto M. 1994. The fission yeast mating pheromone P-factor: its molecular structure, gene structure, and ability to induce gene expression and G1 arrest in the mating partner. Genes Dev 8:328–38. doi:10.1101/gad.8.3.328

Iwao Y. 2014. Egg Activation in Polyspermy: Its Molecular Mechanisms and Evolution in VertebratesSexual Reproduction in Animals and Plants. Tokyo: Springer Japan. pp. 171–180. doi:10.1007/978-4-431-54589-7_15

Kemp HA, Sprague GF. 2003. Far3 and five interacting proteins prevent premature recovery from pheromone arrest in the budding yeast Saccharomyces cerevisiae. Mol Cell Biol 23:1750–63. doi:10.1128/mcb.23.5.1750-1763.2003

Kjaerulff S, Andersen NR, Borup MT, Nielsen O. 2007. Cdk phosphorylation of the Ste11 transcription factor constrains differentiation-specific transcription to G1. Genes Dev 21:347–59. doi:10.1101/gad.407107

Kohli J, Hottinger H, Munz P, Strauss A, Thuriaux P. 1977. Genetic Mapping in Schizosaccharomyces pombe by Mitotic and Meiotic Analysis and Induced Haploidization. Genetics 87:471–89.

Kunitomo H, Higuchi T, Iino Y, Yamamoto M. 2000. A zinc-finger protein, Rst2p, regulates transcription of the fission yeast ste11(+) gene, which encodes a pivotal transcription factor for sexual development. Mol Biol Cell 11:3205–17. doi:10.1091/mbc.11.9.3205

Legland D, Arganda-Carreras I, Andrey P. 2016. MorphoLibJ: integrated library and plugins for mathematical morphology with ImageJ. Bioinformatics 32:3532–3534. doi:10.1093/bioinformatics/btw413

Li P, McLeod M. 1996. Molecular mimicry in development: identification of ste11+ as a substrate and mei3+ as a pseudosubstrate inhibitor of ran1+ kinase. Cell 87:869–80. doi:10.1016/s0092-8674(00)81994-7

Luporini P, Pedrini B, Alimenti C, Vallesi A. 2016. Revisiting fifty years of research on pheromone signaling in ciliates. Eur J Protistol 55:26–38. doi:10.1016/j.ejop.2016.04.006

Martin SG. 2019. Molecular mechanisms of chemotropism and cell fusion in unicellular fungi. J Cell Sci 132. doi:10.1242/JCS.230706

Martin SG, Arkowitz RA. 2014. Cell polarization in budding and fission yeasts. FEMS Microbiol Rev 38:228–253. doi:10.1111/1574-6976.12055

Mata J, Bähler J. 2006. Global roles of Ste11p, cell type, and pheromone in the control of gene expression during early sexual differentiation in fission yeast. Proc Natl Acad Sci U S A 103:15517–22. doi:10.1073/pnas.0603403103

Mata J, Lyne R, Burns G, Bähler J. 2002. The transcriptional program of meiosis and sporulation in fission yeast. Nat Genet 32:143–147. doi:10.1038/ng951

Maundrell K. 1993. Thiamine-repressible expression vectors pREP and pRIP for fission yeast. Gene 123:127–130. doi:10.1016/0378-1119(93)90551-D

McLeod M, Beach D. 1988. A specific inhibitor of the ran1+ protein kinase regulates entry into meiosis in Schizosaccharomyces pombe. Nature 332:509–14. doi:10.1038/332509a0

McLeod M, Stein M, Beach D. 1987. The product of the mei3+ gene, expressed under control of the mating-type locus, induces meiosis and sporulation in fission yeast. EMBO J 6:729–36.

Meister P, Taddei A, Ponti A, Baldacci G, Gasser SM. 2007. Replication foci dynamics: replication patterns are modulated by S-phase checkpoint kinases in fission yeast. EMBO J 26:1315–26. doi:10.1038/sj.emboj.7601538

Merlini L, Khalili B, Bendezú FO, Hurwitz D, Vincenzetti V, Vavylonis D, Martin SG. 2016. Local Pheromone Release from Dynamic Polarity Sites Underlies Cell-Cell Pairing during Yeast Mating. Curr Biol 26:1117–25. doi:10.1016/j.cub.2016.02.064

Merlini L, Khalili B, Dudin O, Michon L, Vincenzetti V, Martin SG. 2018. Inhibition of Ras activity coordinates cell fusion with cell–cell contact during yeast mating. J Cell Biol 217:1467–1483. doi:10.1083/jcb.201708195

Murakami-Tonami Y, Yamada-Namikawa C, Tochigi A, Hasegawa N, Kojima H, Kunimatsu M, Nakanishi M, Murakami H. 2007. Mei4p coordinates the onset of meiosis I by regulating cdc25+ in fission yeast. Proc Natl Acad Sci U S A 104:14688–93. doi:10.1073/pnas.0702906104

Navarro FJ, Chakravarty P, Nurse P. 2017. Phosphorylation of the RNA-binding protein Zfs1 modulates sexual differentiation in fission yeast. J Cell Sci 130:4144–4154. doi:10.1242/jcs.208066

Otsubo Y, Yamashita A, Ohno H, Yamamoto M. 2014. S. pombe TORC1 activates the ubiquitin-proteasomal degradation of the meiotic regulator Mei2 in cooperation with Pat1 kinase. J Cell Sci 127:2639–46. doi:10.1242/jcs.135517

Peter M, Herskowitz I. 1994. Direct inhibition of the yeast cyclin-dependent kinase Cdc28-Cln by Far1. Science (80-) 265:1228–31. doi:10.1126/science.8066461

Petersen J, Weilguny D, Egel R, Nielsen O. 1995. Characterization of fus1 of Schizosaccharomyces pombe: a developmentally controlled function needed for conjugation. Mol Cell Biol 15:3697–707.

Rothbauer U, Zolghadr K, Tillib S, Nowak D, Schermelleh L, Gahl A, Backmann N, Conrath K, Muyldermans S, Cardoso MC, Leonhardt H. 2006. Targeting and tracing antigens in live cells with fluorescent nanobodies. Nat Methods 3:887–889. doi:10.1038/nmeth953

Saito TT, Tougan T, Kasama T, Okuzaki D, Nojima H. 2004. Mcp7, a meiosis-specific coiled-coil protein of fission yeast, associates with Meu13 and is required for meiotic recombination. Nucleic Acids Res 32:3325–39. doi:10.1093/nar/gkh654

Schindelin J, Arganda-Carreras I, Frise E, Kaynig V, Longair M, Pietzsch T, Preibisch S, Rueden C, Saalfeld S, Schmid B, Tinevez J-Y, White DJ, Hartenstein V, Eliceiri K, Tomancak P, Cardona A. 2012. Fiji: an open-source platform for biological-image analysis. Nat Methods 9:676–682. doi:10.1038/nmeth.2019

Shaner NC, Campbell RE, Steinbach PA, Giepmans BNG, Palmer AE, Tsien RY. 2004. Improved monomeric red, orange and yellow fluorescent proteins derived from Discosoma sp. red fluorescent protein. Nat Biotechnol 22:1567–1572. doi:10.1038/nbt1037

Shimada M, Yamada-Namikawa C, Murakami-Tonami Y, Yoshida T, Nakanishi M, Urano T, Murakami H. 2008. Cdc2p controls the forkhead transcription factor Fkh2p by phosphorylation during sexual differentiation in fission yeast. EMBO J 27:132–42. doi:10.1038/sj.emboj.7601949

Snook RR, Hosken DJ, Karr TL. 2011. The biology and evolution of polyspermy: insights from cellular and functional studies of sperm and centrosomal behavior in the fertilized egg. Reproduction 142:779–92. doi:10.1530/REP-11-0255

Stelling J, Sauer U, Szallasi Z, Doyle FJ, Doyle J. 2004. Robustness of cellular functions. Cell 118:675–85. doi:10.1016/j.cell.2004.09.008

Stern B, Nurse P. 1998. Cyclin B proteolysis and the cyclin-dependent kinase inhibitor rum1p are required for pheromone-induced G1 arrest in fission yeast. Mol Biol Cell 9:1309–1321.

Stern B, Nurse P. 1997. Fission yeast pheromone blocks S-phase by inhibiting the G1 cyclin B-p34cdc2 kinase. EMBO J 16:534–44. doi:10.1093/emboj/16.3.534

Strickfaden SC, Winters MJ, Ben-Ari G, Lamson RE, Tyers M, Pryciak PM. 2007. A Mechanism for Cell-Cycle Regulation of MAP Kinase Signaling in a Yeast Differentiation Pathway. Cell 128:519–531. doi:10.1016/J.CELL.2006.12.032

Sugimoto a, Iino Y, Maeda T, Watanabe Y, Yamamoto M. 1991. Schizosaccharomyces pombe ste11+ encodes a transcription factor with an HMG motif that is a critical regulator of sexual development. Genes Dev 5:1990–9. doi:10.1101/gad.5.11.1990

Takeda T, Toda T, Kominami K, Kohnosu A, Yanagida M, Jones N. 1995. Schizosaccharomyces pombe atf1+ encodes a transcription factor required for sexual development and entry into stationary phase. EMBO J 14:6193–208.

Tanaka K, Davey J, Imai Y, Yamamoto M. 1993. Schizosaccharomyces pombe map3+ encodes the putative M-factor receptor. Mol Cell Biol 13:80–8.

Vještica A, Marek M, Nkosi PJ, Merlini L, Liu G, Bérard M, Billault-Chaumartin I, Martin SG. 2020. A toolbox of stable integration vectors in the fission yeast Schizosaccharomyces pombe. J Cell Sci 133. doi:10.1242/JCS.240754

Vjestica A, Merlini L, Dudin O, Bendezu FO, Martin SG. 2016. Microscopy of Fission Yeast Sexual Lifecycle. J Vis Exp e53801–e53801. doi:10.3791/53801

Vještica A, Merlini L, Nkosi PJ, Martin SG. 2018. Gamete fusion triggers bipartite transcription factor assembly to block re-fertilization. Nature 560:397–400. doi:10.1038/s41586-018-0407-5

Wallen RM, Perlin MH. 2018. An Overview of the Function and Maintenance of Sexual Reproduction in Dikaryotic Fungi. Front Microbiol 9:1–24. doi:10.3389/fmicb.2018.00503

Watanabe Y, Yamamoto M. 1994. S. pombe mei2+ encodes an RNA-binding protein essential for premeiotic DNA synthesis and meiosis I, which cooperates with a novel RNA species meiRNA. Cell 78:487–498. doi:10.1016/0092-8674(94)90426-X

Watanabe Y, Yokobayashi S, Yamamoto M, Nurse P. 2001. Pre-meiotic S phase is linked to reductional chromosome segregation and recombination. Nature 409:359–363. doi:10.1038/35053103

Willer M, Hoffmann L, Styrkársdóttir U, Egel R, Davey J, Nielsen O. 1995. Two-step activation of meiosis by the mat1 locus in Schizosaccharomyces pombe. Mol Cell Biol 15:4964–4970.

Win TZ, Gachet Y, Mulvihill DP, May KM, Hyams JS. 2001. Two type V myosins with non-overlapping functions in the fission yeast Schizosaccharomyces pombe: Myo52 is concerned with growth polarity and cytokinesis, Myo51 is a component of the cytokinetic actin ring. J Cell Sci 114:69–79.

Wong JL, Wessel GM. 2005. Defending the Zygote: Search for the Ancestral Animal Block to Polyspermy. Curr Top Dev Biol 72:1–151. doi:10.1016/S0070-2153(05)72001-9

